# Trehalose dimycolate inhibits phagosome maturation and promotes intracellular *M. tuberculosis* growth via noncanonical SNARE interaction

**DOI:** 10.1101/2024.12.16.627577

**Authors:** Carolina Santamaria, Kyle J. Biegas, Pamelia N. Lim, Jessica Cabral, Christi Y. Kim, James R. Lee, Ishani V. Gaidhane, Casey Papson, Kyla Gomard-Henshaw, Alissa C. Rothchild, Benjamin M. Swarts, M. Sloan Siegrist

## Abstract

Mycobacterial cell envelopes are rich in unusual lipids and glycans that play key roles during infection and vaccination. The most abundant envelope glycolipid is trehalose dimycolate (TDM). TDM compromises the host response to mycobacterial species via multiple mechanisms, including inhibition of phagosome maturation. The molecular mechanism by which TDM inhibits phagosome maturation has been elusive. We find that a clickable, photoaffinity TDM probe recapitulates key phenotypes of native TDM in macrophage host cells and binds several host SNARE proteins, including VTI1B, STX8, and VAMP2. VTI1B and STX8 normally promote endosome fusion by forming a complex with VAMP8. However, in the presence of *Mycobacterium tuberculosis*, VTI1B and STX8 complex with VAMP2, which in turn decreases VAMP8 binding. VAMP2 acts together with mycolate structure to inhibit phagosome maturation and promotes intracellular *M. tuberculosis* replication. Thus one mechanism by which TDM constrains the innate immune response to *M. tuberculosis* is via non-canonical SNARE complexation.

**Significance Statement:** Glycolipids from the *Mycobacterium tuberculosis* cell envelope, particularly trehalose dimycolate (TDM), play major roles in subverting the immune response to this intracellular pathogen. How subversion occurs is often unclear because glycans and lipids are technically challenging to study in cells. We discovered that a TDM-mimicking chemical probe interacts with three host SNARE proteins, including two that regulate endosome fusion and one that does not. The presence of TDM or *M. tuberculosis* triggers abnormal binding of these SNAREs, which in turn inhibits the fusion of *M. tuberculosis*-containing phagosomes with lysosomes and promotes *M. tuberculosis* replication. Our work provides an unusual example of a bacterial pathogen restricting the immune response via glycolipid-SNARE interactions.

## Introduction

The cell envelope of mycobacteria shapes the innate immune response to these organisms. Surrounding the cytoplasm, the mycobacterial envelope consists of a peptidoglycan cell wall, an arabinogalactan layer, and a distinctive outer “myco” membrane (Figure 1A) that is unique to the *Corynebacterineae* suborder. The inner and outer leaflets of the mycomembrane are respectively composed of mycolic acids that are covalently attached to arabinogalactan and intercalated glycolipids (1). The most abundant envelope glycolipid, trehalose dimycolate (TDM), has various critical roles in *Mycobacterium tuberculosis* pathogenesis (2). For example, TDM engages several host receptors, including Mincle (Macrophage inducible Ca^2+^-dependent lectin receptor), to trigger a pro-inflammatory cytokine response and granuloma formation (3–5). TDM can also impair the host response and promote intracellular mycobacterial survival by inhibiting phagosome maturation and acidification in host cell macrophages (6, 7).

**Figure 1.**
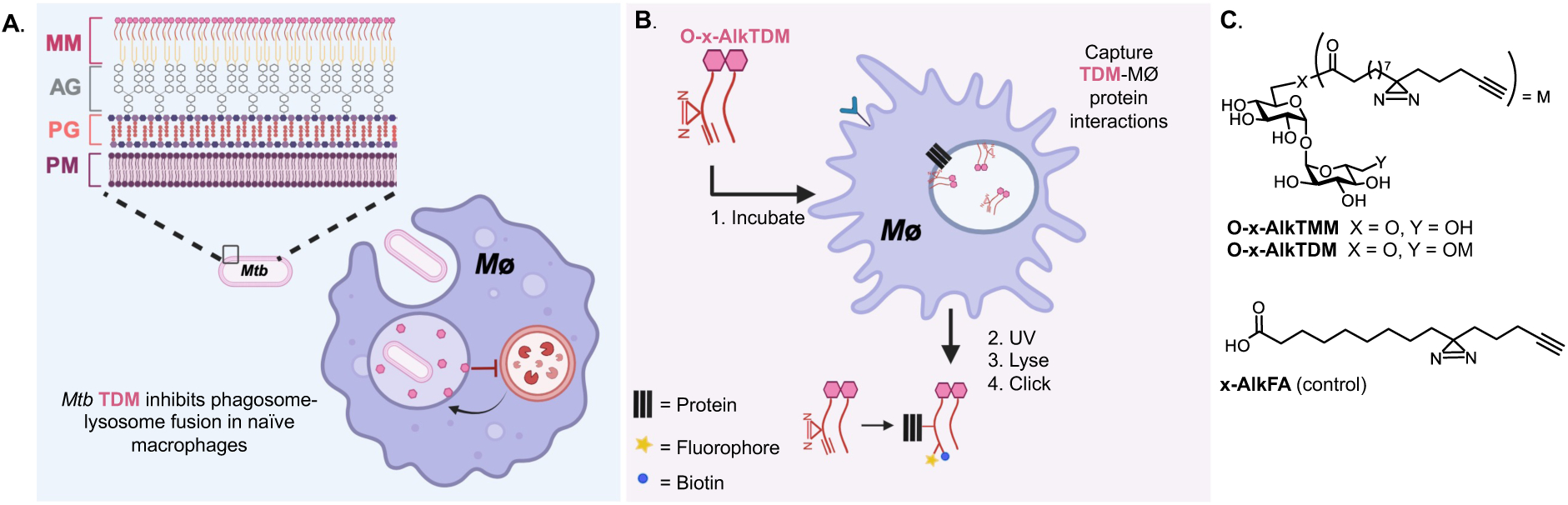
**Chemical probes can reveal interactions between (myco)bacterial lipids and host proteins. (**A) The glycolipid trehalose dimycolate (TDM) in the outer mycomembrane (MM) of *M. tuberculosis* (*Mtb*) inhibits fusion of lysosomes with *Mtb*-containing phagosomes. Other components of the mycobacterial cell envelope include arabinogalactan (AG), peptidoglycan (PG), and the inner plasma membrane (PM). (B) Workflow for chemical proteomics using the TDM-mimicking probe O-x-AlkTDM. (C) Structures of O-x-AlkTDM and control probes O-x-AlkTMM and x-AlkFA.

During normal phagosome maturation, a microbe-containing phagosome fuses with endosomes and eventually lysosomes. Subsequent acidification of the phagolysosome activates enzymes that destroy the microbe, in turn enabling antigen processing and presentation on the phagocytic cell surface that can initiate adaptive immune responses (8, 9). Inhibition of phagosome maturation is a well-documented strategy for immune evasion across bacterial, protozoan, and fungal species, first observed in Mycobacterium tuberculosis and the vaccine M. bovis BCG in 1971 (10). In addition to TDM, mycobacterial species use secreted protein effectors and other envelope lipids to inhibit phagosome maturation (11). To date, the reported mechanisms by which TDM or any envelope lipid interferes with normal phagosome maturation are indirect, e.g., inhibition of host signaling pathways (6), and/or speculative, e.g., modulation of phagosome proteome (12–15) and in vitro inhibition of model liposome fusion (16). Envelope lipids can be released by BCG and the aquatic pathogen M. marinum during infection and “spread” throughout host membranes (17–20). While their physical location in host cells suggests the potential for these lipids to directly interfere with fusion events that mediate phagosome maturation, such mechanisms have not been reported in live cells.

One strategy by which other intracellular pathogens hijack phagosome maturation is via host SNAREs (Soluble N-ethylmaleimide-Sensitive Factor Attachment Proteins Receptor), membrane-bound mediators of fusion between host vesicles and organelles (21, 22). Pathogen interference with the expression and/or function of SNAREs can create a more hospitable intracellular environment by controlling vesicle traffic to and from the phagosome (23, 24). The BCG vaccine alters SNARE recruitment to the phagosome (13, 14, 12), although the downstream consequences on host-BCG interactions have not been demonstrated nor have similar findings yet been reported for pathogenic species such as *M. tuberculosis*. Additionally, while other microbes secrete proteins that mimic or modify the SNAREs, similar-acting mycobacterial proteins have not been identified. Instead, mycobacterial envelope glycolipids like mannose-capped lipoarabinomannan (ManLAM) and its biosynthetic precursor phosphatidylinositol mannoside (PIM) can modulate SNARE recruitment to the phagosome, albeit in ways that are indirect (14) or not yet characterized (12).

While mycobacterial envelope lipids and glycans influence numerous interactions with the host, significant gaps in mechanistic insight persist due to the technical challenges of tracking and manipulating these components within cells. We focused on TDM, an abundant envelope glycolipid that contributes to the pathophysiology of both tuberculosis infection (2, 25) and BCG vaccination (26) in addition to inhibiting phagosome maturation. Here we synthesize and deploy a clickable, photocrosslinking TDM probe to identify interacting proteins in host cell macrophages (Figure 1B). Similar to native TDM, the probe stimulates a pro-inflammatory cytokine response, inhibits phagosome maturation, and interacts with Mincle. Chemical proteomics followed by co-immunoprecipitation experiments revealed that the TDM probe interacts with host SNAREs. Three of the SNAREs that we identified–VTI1B, STX7, and STX8– comprise a late endosome complex that normally binds VAMP7 or VAMP8 to mediate fusion with other endosomes or with lysosomes (21, 27, 28). The fourth SNARE that we identified, VAMP2, can associate with endosomes and lysosomes but it is not known to bind VTI1B, STX7, or STX8 (21, 29, 30). We find that VAMP2 forms a non-canonical complex with VTI1B and STX8 specifically in the presence of native TDM or *M. tuberculosis*. During macrophage infection, VAMP2 inhibits VAMP8 complexation with VTI1B and STX8, acts with mycolic acids to inhibit phagosome maturation, and promotes *M. tuberculosis* replication. Thus, TDM drives non-canonical SNARE complexation to benefit intracellular *M. tuberculosis* survival (7, 31–33).

## Results

### Mycobacterial Glycolipid Probe O-x-AlkTDM Recapitulates Key Aspects of TDM Activity and Binding in Macrophages

How TDM inhibits phagosome maturation is unknown. We reasoned that identification of macrophage proteins that interact with TDM could provide clues to the mechanism of TDM’s inhibition. However, glycolipid-protein interactions are transient and generally weaker than protein-protein interactions, and therefore less amenable to analysis using traditional biochemical approaches. We previously reported a clickable, photocrosslinking trehalose monomycolate (TMM) probe, which is converted by live mycobacteria into TDM, and used this technology to identify TDM-interacting proteins in the model organism *M. smegmatis* (34). To identify TDM-interacting proteins in host cells, we synthesized O-x-AlkTDM, a direct mimic of TDM bearing an alkyne and a diazirine with linear, truncated lipid chains (Figure 1C). The diazirine enables crosslinking upon ultraviolet light irradiation while the alkyne enables copper-catalyzed alkyne-azide cycloaddition (CuAAC) to attach azide-bearing detection moieties. The structure of O-x-AlkTDM was inspired by the well-established, TDM-mimicking adjuvant trehalose-6,6-dibehenate (TDB), which elicits a macrophage response similar to native TDM despite its simplified lipid chains (35). The goal of the compound design was to create a probe that enables covalent capture and pull-down assays while recapitulating the phenotypes of native TDM or TDB adjuvant in host cells (Figure 1B). O-x-AlkTDM was chemically synthesized using our established strategy and its key functionalities were validated in a photocrosslinking experiment with bovine serum albumin (Supporting Information (SI), Figures S1–S4).

We first tested whether the O-x-AlkTDM probe stimulates a pro-inflammatory cytokine response. After incubating immortalized bone marrow-derived macrophages (iBMDMs) (37) from C57BL/6 mice with O-x-AlkTDM, we observed that the probe triggered a cytokine response similar to that of TDB, *i.e*., dose-dependent TNF-alpha and IL-6 production (Figures 2A-B, S5A-B). We next asked whether O-x-AlkTDM inhibits phagosome maturation. We coated fluorescent latex beads with native TDM, TDB, or O-x-AlkTDM, incubated with macrophages differentiated from the human monocyte cell line THP-1, and quantitated the proportion of beads that were spatially coincident with LysoTracker, a dye that stains acidic compartments. All three molecules decreased spatial coincidence of the beads with LysoTracker relative to bovine serum albumin (BSA)-coated control beads, although native TDM did so more potently than either TDB or O-x-AlkTDM (Figure 2C-D and SI, Figure S5C). We then localized O-x-AlkTDM within iBMDMs after CuAAC with Picolyl Alexa 488 Azide. Consistent with previous observations that mycobacterial envelope lipids, including TDM, can traffic throughout the endocytic network (18, 38), O-x-AlkTDM appeared to form cytoplasmic puncta (SI, Figure S6). Finally, we determined whether

**Figure 2.**
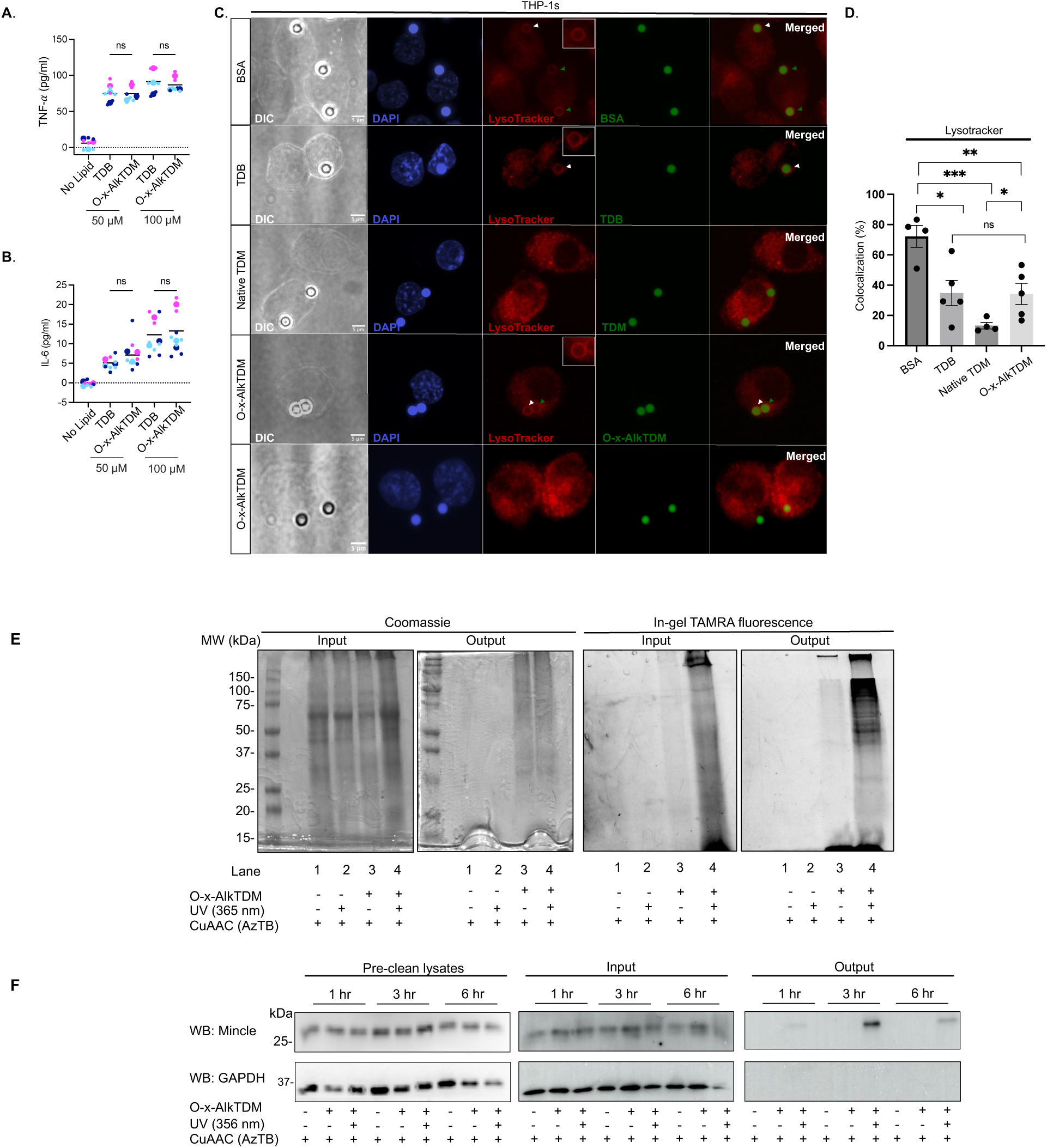
Mycobacterial glycolipid probe O-x-AlkTDM recapitulates key aspects of TDM activity and binding in macrophages. ELISA detection of pro-inflammatory cytokines (A) TNF-alpha and (B) IL-6 after 6 hr incubation of immortalized bone marrow-derived macrophages (iBMDMs) from C57BL/6 mice with O-x-AlkTDM or commercial TDM mimic trehalose dibehenate (TDB). For (A-B) each color in the SuperPlots (36) represents an independent biological replicate. Smaller symbols represent technical replicates, and larger symbols are the means of the technical replicates per experiment. Statistical significance determined using a two-way ANOVA with a Šídák’s multiple comparisons test for post hoc analysis. Data obtained from three independent experiments. (C-D) Lysosome fusion of bead-containing phagosomes. THP-1 cells were incubated with BSA-, TDB-, TDM-, or O-x-alkTDM-coated fluorescent beads (green), followed by LysoTracker staining to assess acidification. Green and white arrows indicate LysoTracker-localized beads; white arrows highlight beads that have been magnified for enhanced visualization. Representative images shown in (C) and (SI, **Figure S5C**). In (D), spatial coincidence between beads and LysoTracker was quantitated from 3-5 independent biological replicates. Images were blinded prior to analysis. White arrows indicate LysoTracker-localized beads that have been magnified for enhanced visualization, while green arrows highlight additional Lysotracker staining present within the image. (E-F) O-x-AlkTDM-mediated affinity enrichment of host interacting proteins. iBMDMs were incubated with O-x-AlkTDM (100 µM), UV-irradiated and lysed. Lysates were reacted with TAMRA Biotin Azide (AzTB) via copper-catalyzed alkyne-azide cycloaddition (CuAAC) and analyzed by Coomassie and in-gel TAMRA fluorescence (input). Clicked samples were incubated with NeutrAvidin agarose beads (output) to evaluate global enrichment of proteins (E) and specific enrichment of the known TDM receptor Mincle (F). Representative data shown for three independent biological replicates (E-F). Additional controls for (E-F) in SI, **Figure S5D**.

O-x-AlkTDM interacts with Mincle, a known receptor for TDM (3, 4, 6, 5). iBMDMs treated with O-x-AlkTDM were irradiated with UV and subjected to CuAAC with TAMRA Biotin Azide (AzTB; Figure 2E). After affinity enrichment and SDS-PAGE, we detected O-x-AlkTDM-interacting proteins via in-gel fluorescence to visualize TAMRA labeling. Immunoblotting then revealed time-dependent interactions between O-x-AlkTDM and Mincle, which peaked at 3 hours (Figure 2F). These findings demonstrate that the O-x-AlkTDM probe can both elicit and interfere with physiologically-relevant innate immune responses and can report a known host protein interaction.

### Macrophage SNAREs Interact with O-x-AlkTDM

We next investigated whether O-x-AlkTDM could reveal new interactions with host proteins. To prepare samples for chemical proteomics, we again treated iBMDMs with the probe, UV-irradiated, clicked with AzTB, and affinity enriched prior to LC-MS/MS. Over-Representation Analysis (39) showed significant enrichment of four SNARE proteins in the presence of O-x-AlkTDM: Syntaxin 8 (STX8), Syntaxin 7 (STX7), Vesicle Associated Membrane Protein 2 (VAMP2), and Vesicle Transport through Interaction with t-SNAREs 1B (VTI1B; Figure 3A). STX8, STX7, and VTI1B were previously reported to form an endosome complex that can bind to either VAMP7 or VAMP8 to mediate homotypic fusion with other endosomes or heterotypic fusion with lysosomes (27, 28). STX8 has also been shown to reside on both latex bead- and BCG-containing phagosomes (14). VAMP2 has primarily been implicated in vesicle exocytosis and in complexes with SNAP25 and STX1A (40); it has not previously been linked to STX8, STX7, or VTI1B (27–30, 41). Co-immunoprecipitation experiments confirmed interactions between O-x-AlkTDM and STX8, VTI1B, and VAMP2 in both iBMDMs and the human cell line THP-1 (Figures 3B-C; we were unable to detect STX7 even from protein lysates).

**Figure 3.**
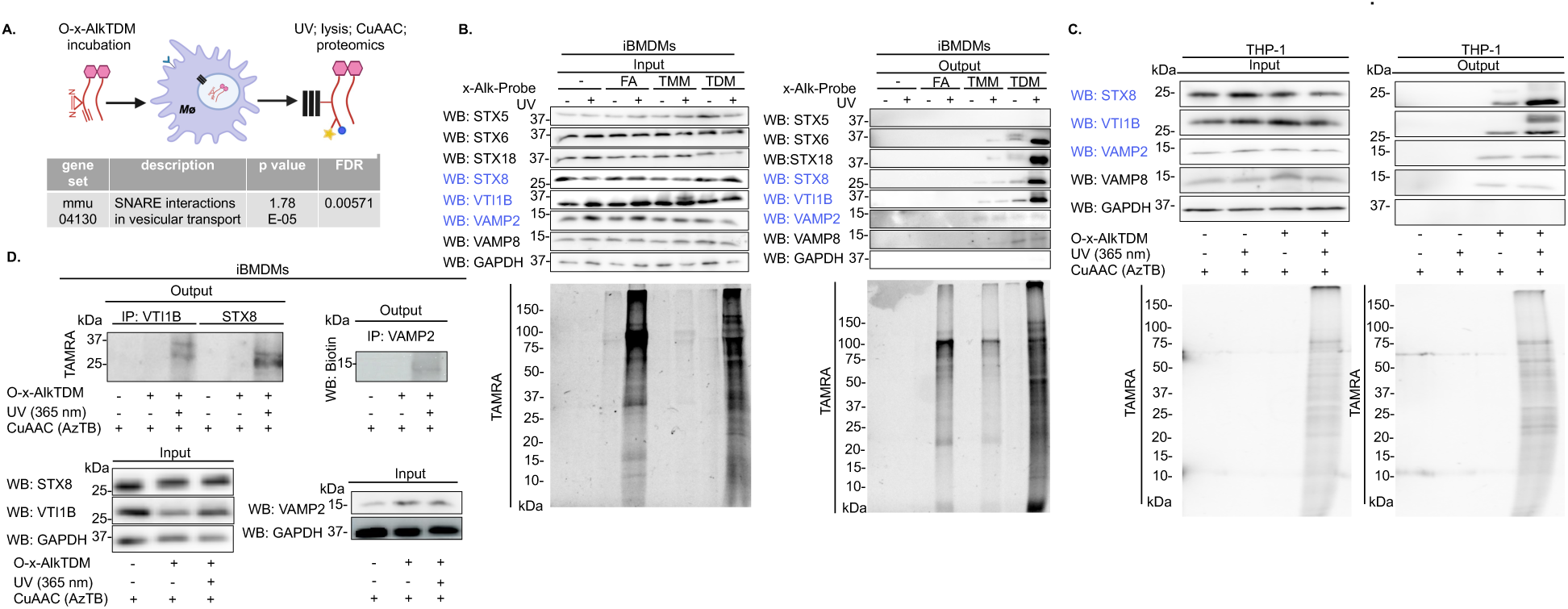
Macrophage SNAREs interact with O-x-AlkTDM. (A) iBMDMs were incubated +/- O-x-alkTDM for 6 hrs; exposed to UV or not; lysed; and subjected to CuAAC with AzTB as in Figures 1B and **2E-F**. O-x-AlkTDM-interacting proteins were then enriched by NeutrAvidin beads; digested with trypsin; and analyzed by LC-MS/MS. Over-Representation Analysis (ORA) performed via WEBGestalt (39). Cut-off set at p<0.05 and false discovery rate (FDR)<0.25. Enrichment is relative to no probe samples across four independent biological replicates for each sample. iBMDMs (B) or THP-1s (C) treated 6 hrs +/- O-x-AlkTDM and processed as in (A) and Figure 2E-F. Interacting proteins before (input) and after affinity enrichment (output) with NeutrAvidin agarose beads detected by in-gel fluorescence (TAMRA) or by immunoblotting with the indicated anti-SNARE antibodies. SNAREs identified by proteomics (A) highlighted in blue. Other SNAREs and housekeeping protein GAPDH in black. In (B), iBMDMs were also treated with control x-Alk-probes that lack either a lipid chain (TMM) or trehalose (FA; for structures, see Figure 1C). Double bands for STX8 and VTI1B blots may be SNAREs interacting with one or two molecules of probe (see **SI, Figure S7**). Representative data shown for 2-5 independent biological replicates. (D) iBMDMs treated as in (B) but immunoprecipitated using the indicated anti-SNARE antibody. The presence of O-x-AlkTDM was confirmed by in-gel fluorescence (TAMRA) or immunoblotting with anti-biotin. There were no obvious changes in protein expression in samples prior to immunoprecipitation (input).

As expected, most O-x-AlkTDM-SNARE interactions were promoted by UV. However, VAMP2 detection after O-x-AlkTDM affinity purification was consistently independent of UV, in both iBMDMs and THP-1s, despite the UV dependence for the reciprocal experiment (O-x-AlkTDM detection before and after VAMP2 immunoprecipitation; Figure 3D). While we do not yet understand why O-x-AlkTDM-VAMP2 is detectable without cross-linking, we hypothesize that this interaction is more stable than O-x-AlkTDM-STX8 or O-x-AlkTDM-VTI1B. We frequently observed multiple bands in pulldowns from the O-x-AlkTDM-treated samples, particularly for VTI1B and STX8 (SI, Figure S7). While not definitive, these data suggest that more than one molecule of O-x-AlkTDM may be able to bind to these proteins. Taken together, our data suggest that O-x-AlkTDM interacts with host SNARE proteins STX8, VTI1B, and VAMP2.

### Specificity of Glycolipid-SNARE Interactions

Structural variability in glycan head group and/or mycolic acids can modulate the host cell response to TDM (4, 42–44). Therefore, we sought to understand the specificity of O-x-AlkTDM-SNARE interactions. To investigate the role of glycolipid structure, we incubated iBMDMs with two additional probes: O-x-AlkTMM (34), which retains the trehalose sugar and one lipid chain, and x-AlkFA, which contains a single lipid chain and lacks the trehalose head group (Figure 1C). Like TDM, TMM is a natural constituent of the mycobacterial cell envelope and a reported ligand for Mincle (5, 44). Following UV irradiation, CuAAC with AzTB, and affinity purification, we found that all three probes produced UV- and click-dependent bands. However, x-AlkFA failed to pull down STX8 or VTI1B. While O-x-AlkTMM pulled down both proteins, it did so less efficiently than O-x-AlkTDM (Figure 3B). These findings suggest that glycolipid-SNARE interactions require a glycan head group and are enhanced by the presence of two lipid chains.

To investigate glycolipid-SNARE specificity on the protein side of the interaction, we immunoblotted samples that had been immunoprecipitated for O-x-AlkTDM/TMM-interacting proteins with antibodies to other SNAREs. The repertoire of SNAREs that O-x-AlkTDM/TMM could pull down extended beyond the ones that we identified by proteomics, and included STX6, STX18, and VAMP8 (Figure 3B-C). VAMP8 mediates endosomal fusion, in complex with VTI1B, STX7, and STX8 (27, 28). As with VAMP2, the O-x-AlkTDM interaction with VAMP8 was not dependent on cross-linking, suggesting that it may be more stable than O-x-AlkTDM-STX6, - STX8, -STX18, or -VTI1B. STX6 and STX18 are respectively involved in trans-Golgi-endosome and ER-Golgi trafficking (22, 21). STX6 is present on latex bead-containing phagosomes but excluded from BCG-containing phagosomes via an indirect mechanism that depends on the mycobacterial glycolipid mannose-capped lipoarabinomannan (ManLAM) (14). Not all of the proteins that we tested pulled down with O-x-AlkTDM; we did not detect interactions between the probe and STX5, a SNARE that mediates trafficking from phagosomes to the trans-Golgi network, nor to GAPDH, a cytosolic enzyme (21, 45) (Figures 3B). We conclude that O-x-AlkTDM selectively interacts with a subset of SNAREs.

### *Mycobacterium tuberculosis* TDM Induces Non-Canonical, VAMP2-containing SNARE Complexation in Macrophages

SNARE complexation pulls membranes together, facilitating fusion necessary for processes like phagosome maturation. SNAREs generally interact to form a stable, four-helix bundle, with Qa-, Qb-, and Qc-SNAREs on one membrane and R-SNAREs on the other membrane (21).

SNAREs can sometimes substitute for one another, a property exploited by various intracellular pathogens that disrupt normal complexation via SNARE deubiquitination (46–48) and secreting SNARE-mimicking proteins (49–51). STX7 (Qa-SNARE), VTI1B (Qb-SNARE), and STX8 (Qc-SNARE) on endosomal membranes fuse with VAMP7 or VAMP8 (R-SNAREs) on endosomal and lysosomal membranes *in vitro* (27, 28). VAMP2 is an R-SNARE that can also associate with endosomes and lysosomes but is not known to interact with STX7, VTI1B, or STX8 (21, 27).

We wondered whether TDM-SNARE interactions might induce and/or mark the formation of non-canonical complexes in which VAMP2 substitutes for VAMP8. We first asked whether VAMP2-containing complexes were detectable in the presence of *M. tuberculosis* by immunoprecipitating SNAREs from iBMDMs infected with *M. tuberculosis* mc^2^6206 (H37Rv Δ*panCD* Δ*leuCD* (52)). We found that VTI1B and STX8 interact with each other regardless of infection status (Figure 4A). By contrast, VTI1B and STX8 partner with VAMP2 only in the presence of *M. tuberculosis* and with a concomitant reduction in binding to VAMP8 (Figure 4A).

**Figure 4.**
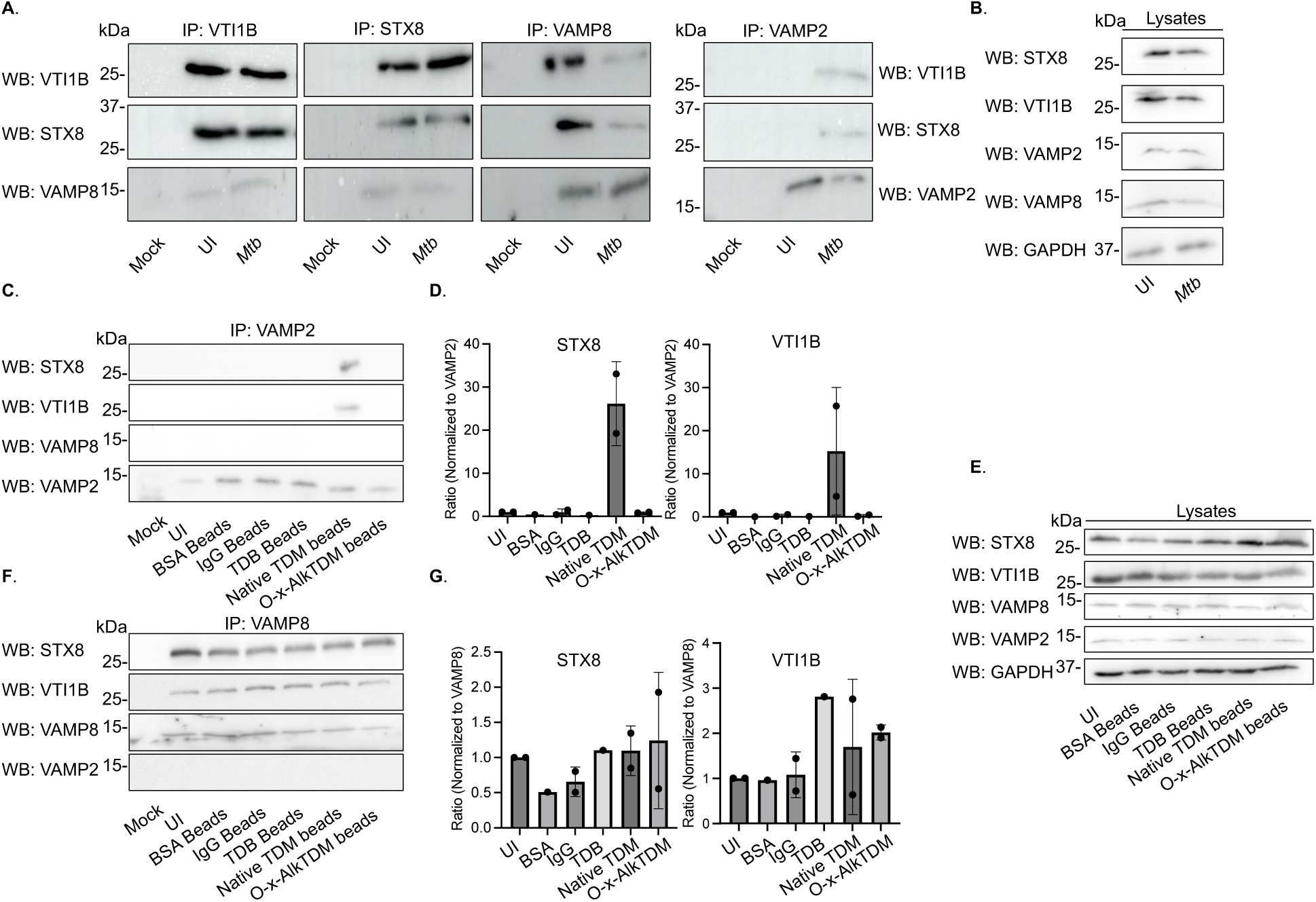
Non-canonical SNARE complexation is triggered by *M. tuberculosis* or native TDM. (A) iBMDMs were infected +/- *M. tuberculosis* mc^2^6206 (*Mtb*; estimated MOI ∼5) then immunoprecipitated and immunoblotted with indicated anti-SNARE antibodies. The VTI1B-STX8 interaction is *M. tuberculosis*-independent, whereas VAMP2 and VAMP8 interactions with VTI1B-STX8 occur preferentially when *M. tuberculosis* is present or absent, respectively. (B) SNARE expression (10 µg/lane) +/- *M. tuberculosis*. iBMDMs infected as in (A) and lysates were immunoblotted with indicated anti-SNARE antibodies. (C-G) iBMDMs were incubated +/- latex beads coated with BSA, IgG, TDB, native TDM, or O-x-AlkTDM then processed as in (A). Native TDM is sufficient to trigger VAMP2 interaction with VTI1B and STX8 (C-D), although a concomitant change in VAMP8 interactions with these proteins was not detected (F-G) in contrast to *M. tuberculosis* (A). (D, G) Quantitation of immunoprecipitations shown in (C) and (G) (and replicates thereof shown in **SI, Figure S8A**), respectively. Mock, isotype control for each primary antibody. UI, uninfected (A-B) or unincubated (C-G). Representative data shown for 2-3 independent biological replicates. (E) SNARE expression (10 µg/lane) +/- latex beads. iBMDMs lysates were immunoblotted with indicated anti-SNARE antibodies prior to immunoprecipitations in (C) and (F).

Protein levels of VAMP2, VAMP8, VTI1B, and STX8 in iBMDMs were similar with or without the pathogen (Figure 4B), consistent with a difference in protein interactions rather than expression. To test whether TDM was sufficient for non-canonical SNARE complexation, we incubated iBMDMs with latex beads coated with BSA, IgG, TDB, O-x-AlkTDM, or native TDM. While VTI1B and STX8 interacted with each other regardless of bead coating, VTI1B and STX8 interacted with VAMP2 only when beads were coated with native TDM (Figure 4C-D). The overall abundance of VAMP2, VAMP8, VTI1B, and STX8 were comparable across bead coatings (Figure 4E). Thus, despite the ability of the O-x-AlkTDM probe to interact with VTI1B, STX8, and VAMP2 (Figure 3), native TDM structure is required for detectable complexation of these proteins. Unlike *M. tuberculosis* infection (Figure 4A-B), TDM-bead incubation did not alter VAMP8 complexation with VTI1B and STX8 (Figure 4F-G). This discrepancy may reflect differential abundance and/or presentation of TDM, *i.e*., bead vs. surface of bacterium, or may indicate that additional *M. tuberculosis* factors are necessary to inhibit normal SNARE complexation.

To test whether non-canonical SNARE complexation triggered by *M. tuberculosis* was dependent on VAMP2, we compared wild-type to *VAMP2* knockout THP-1 cells. As in iBMDMs, *M. tuberculosis* mc^2^6206 at a high estimated multiplicity of infection (MOI ∼5) reduced VAMP8 complexation in wild-type THP-1 cells. However in *VAMP2* knockout THP-1s, VAMP8 interactions with VTI1B and STX8 were similar in the presence or absence of *M. tuberculosis* (Figures 5A-C and SI, Figure S8). We observed a similar phenotype for the fully-virulent parental strain of *M. tuberculosis* H37Rv at high estimated MOI ∼3 (Figure 5D-F). Taken together, our data suggest that *M. tuberculosis* TDM triggers the formation of non-canonical, VAMP2-containing SNARE complexes at the expense of cognate VAMP8-containing complexes.

**Figure 5.**
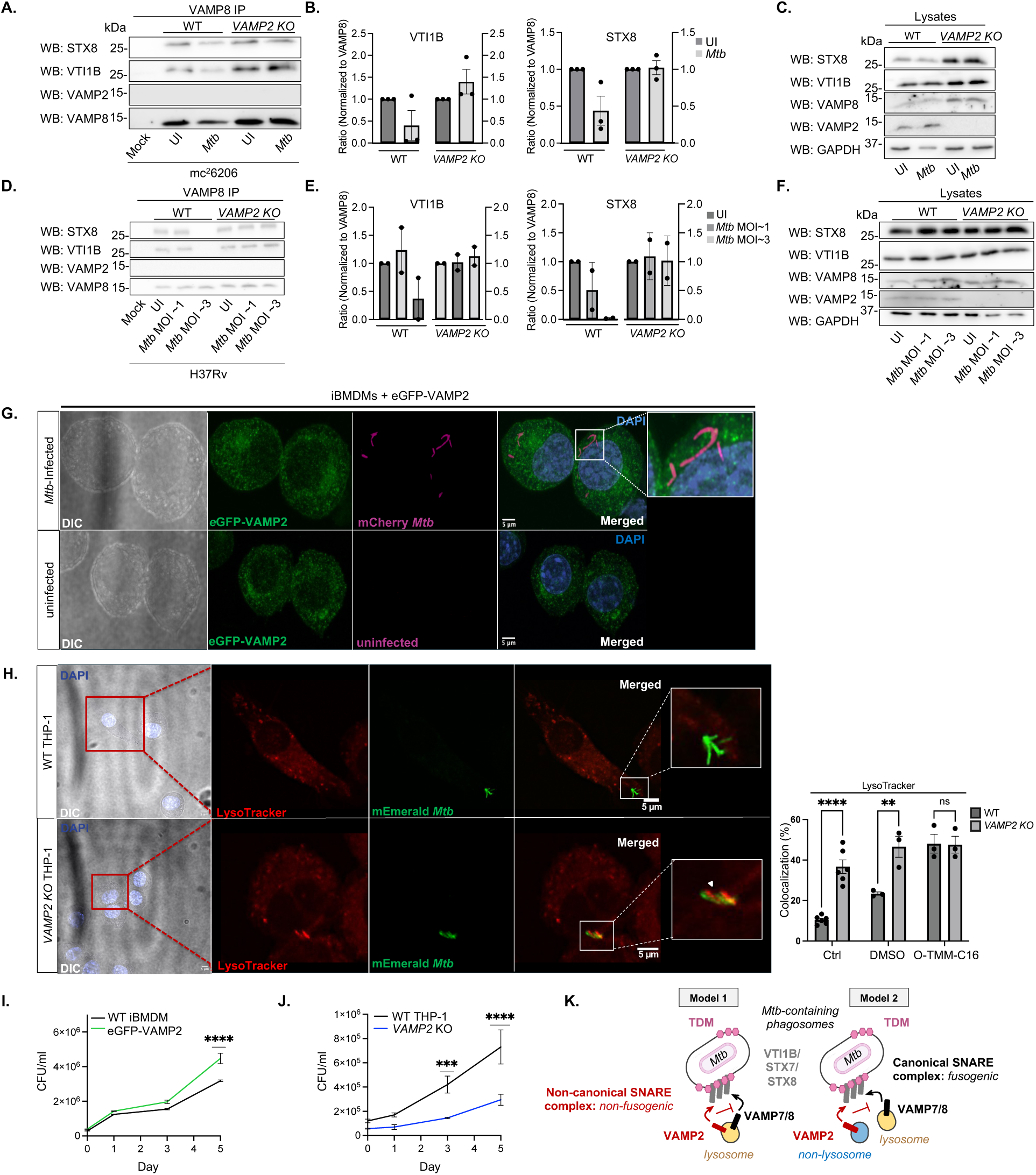
VAMP2 inhibits phagosome acidification and promotes *M. tuberculosis* survival during macrophage infection. Wild-type (WT) and *VAMP2* knockout (KO) THP-1s infected +/- *M. tuberculosis* mc^2^6206 (estimated MOI ∼5) or fully-virulent parent strain H37Rv (estimated MOI ∼1 or 3) and immunoblotted with anti-SNAREs after (A, D) or before (C, F) immunoprecipitation. (B) and (D) show quantitation of immunoprecipitations from (A) and (E), respectively, as well as additional replicates from **SI, Figure S8**. VAMP8 complexation with VTI1B and STX8 in the presence of high MOI *M. tuberculosis* is enhanced in the absence of VAMP2 for both strains. Mock, isotype control for each primary antibody. UI, uninfected. (G) iBMDMs expressing eGFP-VAMP2 were infected +/- mCherry-expressing *M. tuberculosis* (estimated MOI ∼1). Additional images available in **SI, Figure S10**. (H) Loss of macrophage VAMP2 or remodeled *M. tuberculosis* mycolates are non-additively associated with enhanced phagosome acidification. THP-1s were infected with mEmerald-expressing *M. tuberculosis* that were untreated (Ctrl) or pre-treated with DMSO carrier or DMSO plus O-TMM-C16 for ∼5 doublings prior to infection (growth curves of treated *M. tuberculosis* in **SI, Figure S12**). Representative images show spatial coincidence between *M. tuberculosis* and LysoTracker (denoted with white arrow) in THP-1 cells. Blinded quantification, right, from 3-6 independent biological replicates each with n=13-26 individual bacteria. Statistical significance was determined using unpaired Student’s t test. Additional images available in **SI, Figure S12**. (I-J) VAMP2 promotes intracellular *M. tuberculosis* survival. iBMDMs expressing eGFP-VAMP2 or not (I) and wild-type and *VAMP2* knockout THP-1s (J) were infected +/- *M. tuberculosis* mc^2^6206 (estimated MOI ∼1). Macrophages were lysed and plated for colony-forming units at the indicated time points after infection. Representative data shown for 2-3 independent biological replicates. Statistical significance was determined using a two-way ANOVA with a Šídák’s multiple comparisons test for post hoc analysis. (K) Proposed model for TDM-SNARE interactions. TDM released from the mycomembrane of *M. tuberculosis* (Figure 1A) intercalates into macrophage membranes, including those of the phagosome and other organelles and vesicles. TDM directly or indirectly promotes non-canonical, VAMP2-containing SNARE complexation at the expense of canonical, VAMP8-containing SNARE complexation. Fusion of lysosome with the *M. tuberculosis*-containing phagosome is suppressed as in Model 1 or Model 2 (details in text), enabling the pathogen to evade destruction by naive macrophages.

To assess the spatial distribution of VAMP2, we nucleofected iBMDMs with a previously-described plasmid expressing eGFP-VAMP2 (53) (SI, Figure S9). While eGFP-VAMP2 localized near intracellular *M. tuberculosis*, as expected, its subcellular distribution did not obviously change in infected vs. uninfected iBMDMs (Figure 5G and SI, Figure S10A). These data suggest that either *M. tuberculosis* changes the behavior of VAMP2 that is already in the vicinity of the pathogen, or that only a small pool of VAMP2, not detectable with our current methods, is recruited to *M. tuberculosis*-containing phagosomes.

### VAMP2 Promotes *M. tuberculosis* Survival in Macrophages

In unactivated macrophages, *M. tuberculosis* is able to replicate well in part because it delays the fusion of phagosomes with low-pH lysosomes (10, 54–56). Given that VTI1B, STX8, and VAMP8 mediate endosome fusion, and that VAMP8 polymorphisms are associated with human susceptibility to pulmonary tuberculosis (57), we hypothesized that VAMP2-dependent disruption of normal VTI1B-STX8-VAMP8 complexation inhibits phagosome-lysosome fusion and promotes *M. tuberculosis* replication. To test, we first infected THP-1s with mEmerald-expressing *M. tuberculosis* and quantified the proportion of bacteria that were spatially coincident with LysoTracker, a dye that stains acidic compartments. More *M. tuberculosis* were spatially coincident with LysoTracker in *VAMP2* knockout compared to wild-type THP-1-s (Figure 5H and SI, Figures S10B and S11), suggesting that VAMP2 helps *M. tuberculosis* to inhibit phagosome-lysosome fusion.

We next sought to test whether cell envelope mycolates like TDM contribute to VAMP2 promotion of phagosome-lysosome fusion. Like O-x-AlkTDM, TDB has shortened, unbranched lipid chains and stimulates cytokine production and Mincle binding (Figures 1C, 2A-B and SI, Figure S5A-B, (3, 58)) but fails to elicit noncanonical VAMP2 complexation (Figure 4C-D) and restricts phagosome maturation less than native TDM (Figures 2C-D). In the Mycobacteriales animal pathogen *Rhodococcus equi*, TDM with truncated mycolic acids inhibits phagosome-lysosome fusion less than full-length TDM (59), suggesting that lipid structure contributes to *R. equi* TDM function. We reasoned that *M. tuberculosis* with truncated, simplified mycolic acids would therefore enable us to discern whether macrophage VAMP2 acts in concert with *M. tuberculosis* mycolates to inhibit phagosome maturation. To test, we remodeled the mycolic acids of mEmerald-expressing *M. tuberculosis* using our previously-reported (60) trehalose monomycolate (O-TMM-C16) mimic with shortened, unbranched lipid chains and used the remodeled strain to infect macrophages. While treatment with the DMSO carrier alone mildly reduced *M. tuberculosis* growth prior to infection, O-TMM-C16 had no additional effect (SI, Figure S12), suggesting that the mycolic acid remodeling does not obviously impair *M. tuberculosis* fitness prior to infection. In wild-type THP-1s, we found that a higher proportion of O-TMM-C16-treated *M. tuberculosis* were spatially coincident with LysoTracker compared to untreated or DMSO carrier-treated *M. tuberculosis* (Figure 5H). These data are consistent with the findings from *R. equi* (59) and support the notion that mycolic acid structure contributes to *M. tuberculosis* inhibition of phagosome acidification. LysoTracker coincidence increased in *VAMP2* knockout relative to wild-type THP-1s upon infection with untreated or DMSO carrier-treated *M. tuberculosis*, but not with O-TMM-C16-treated *M. tuberculosis*. The non-additive phenotypes suggest that *M. tuberculosis* cell envelope mycolates and macrophage VAMP2 function in the same pathway to impede phagosome maturation.

Finally, to investigate a role for VAMP2 in *M. tuberculosis* replication, we then infected wild-type and eGFP-VAMP2-expressing iBMDMs (which express both eGFP-VAMP2 (53) and VAMP2 (SI, Figure S9)) and wild-type and *VAMP2* knockout THP-1s and plated for colony-forming units at various time points. We found that *M. tuberculosis* replicated significantly better in eGFP-VAMP2-expressing iBMDMs compared to wild-type, and more poorly in VAMP2 knockout THP-1s compared to wild-type (Figures 5I-J). These findings indicate that VAMP2 promotes *M. tuberculosis* replication in addition to inhibiting phagosome acidification.

## Discussion

Lipids and glycans are released from the mycobacterial envelope during infection and affect host cell function. Chief among these bioactive molecules is TDM, a highly abundant envelope glycolipid that induces granuloma formation, stimulates pro-inflammatory cytokine production, and inhibits phagosome maturation (2, 3, 7, 16, 31). Despite the importance of TDM and other envelope components to *M. tuberculosis* infection and *M. bovis* BCG vaccination, the mechanistic details of how TDM subverts the immune response have remained unclear. To facilitate the study of TDM in host cells, we drew on recent advances in chemical techniques to profile glycan- and lipid-protein interactions (34, 61–63). We found that the photocrosslinking, clickable TDM probe O-x-AlkTDM recapitulates key effects of TDM on macrophages and interacts with host SNAREs previously implicated in endosome fusion, including VTI1B, STX8, and potentially STX7. We also discovered that a different O-x-AlkTDM-interacting SNARE, VAMP2, forms a non-canonical complex with VTI1B and STX8 in the presence of native TDM or *M. tuberculosis*, with a reduction in VAMP8 complexation in the latter condition. Many other intracellular pathogens usurp SNARE function; they do so by secreted protein effectors that mimic, degrade, or modify the SNAREs (24, 49–51, 23). To our knowledge, this is the first report of a bacterial lipid interfering with SNARE function.

How does TDM promote VAMP2 interaction with STX8 and VTI1B? O-x-AlkTDM directly interacts with several SNAREs, all of which are predicted to have transmembrane domains. As mycobacterial envelope lipids, including TDM, can traffic throughout the endocytic network (17–20), we hypothesize that TDM intercalates into host membranes and interacts with SNARE transmembrane domains (Figure 5K). Further application of O-x-AlkTDM may enable precise identification of these binding sites, *e.g*., by quantitative crosslinking mass spectrometry.

Model and mammalian cell membranes rigidify in the presence of TDM (64, 65). Many SNAREs localize to ordered membrane regions (66) or, like VAMP2, undergo structural changes that promote complexation (67). The transmembrane domain of VAMP2 in particular is sensitive to lipid bilayer composition (68). More broadly, membrane lipids are well-known to affect SNARE clustering, conformation, and fusion (69, 70). Thus, we posit that TDM-triggered changes to the biophysical milieux of host membranes – alone or in addition to direct interactions with SNARE transmembrane domains – promote complexation of R-SNARE VAMP2 with Q-SNAREs VTI1B, STX8, and STX7 (Figure 5K). In this model, VAMP2 engagement inhibits complexation of the Q-SNAREs with cognate R-SNARE VAMP8. While TDM could promote non-canonical complexation via its effects on VTI1B, STX8, and/or VAMP2, we favor the latter because VAMP2 is known to be impacted by lipid composition (67, 68) and because O-x-AlkTDM interacts especially tightly with VAMP2 compared to VTI1B or STX8 (Figure 3B-C).

O-x-AlkTDM can interact with VTI1B, STX8, and VAMP2 (Figure 3), but, like TDB, seemingly does not trigger non-canonical complexation of these SNAREs like TDM (Figure 4C-D).

Because O-x-AlkTDM and TDB mimic other important aspects of native TDM interaction with macrophages (Figure 2), we speculate that they are similar enough to TDM that they can interact with the same host SNAREs but have structural modifications that weaken binding strength and/or kinetics—either with the SNAREs themselves or with other components of the phagosome membrane—and therefore fail to promote detectable, noncanonical complexation between the SNAREs. While here we took advantage of mycolic acid modifications to test the role of lipid structure in phagosome-lysosome fusion and epistasis with macrophage *VAMP2* (Figure 5H), future iterations of glycolipid probes with more native-like lipid chain structures are high-priority targets for development and downstream structure-activity relationship studies (71, 72).

VTI1B, STX8, and STX7 likely reside on the *M. tuberculosis*-containing phagosome, but the nature of the R-SNARE target membranes is not yet clear. For example, VAMP2 and VAMP8 may both reside on lysosomal membranes (Fig. 5K; Model 1). In this scenario, TDM-triggered engagement of VTI1B, STX8, and STX7 with VAMP2 results in a non-functional complex and stalling of phagosome-lysosome fusion. Alternatively, or additionally, VAMP2 may reside on the membrane of a different vesicle and inhibit the fusion of VAMP8 lysosomes with VTI1B/STX8/STX7-containing phagosomes (Figure 5K; Model 2). High resolution imaging in macrophages, enabled by the clickable tag on O-x-AlkTDM, together with *in vitro* systems with purified organelles, may provide spatial and topological insights into how TDM disrupts vesicle trafficking.

VAMP2 was previously identified as a constituent of the macrophage phagosome proteome that promotes phagocytosis of *E. coli* (73). While we do observe a contribution of VAMP2 to *M. tuberculosis* uptake, the kinetics of replication indicate that VAMP2 additionally promotes intracellular *M. tuberculosis* growth (Figure 5I-J), likely via its effect on phagosome maturation (Figure 5H). Most studies on VAMP2 have focused on its role in secretory vesicles and neuronal exocytosis, raising the possibility that TDM interaction with this SNARE could affect exocytosis from *M. tuberculosis*-infected macrophages, an additional pathway for immune modulation (74). This is especially intriguing given that mycobacterial glycolipids, including TDM, eventually traffic to lysosome-like compartments that are exocytosed (17, 75).

The BCG vaccine does not reliably protect against adult pulmonary tuberculosis, the most common form of the disease. Many groups have developed new recombinant strains, methods of preparation, and modes of administration to enhance the vaccine’s efficacy. BCG, like *M. tuberculosis*, inhibits phagosome maturation (10), which in turn decreases antigen processing, the Th1 response, and efficacy (76–79). Targeting of the TDM-VAMP2 interaction, *e.g*., via transient chemical remodeling of BCG mycolates (Figure 5H), has the potential to enhance antigen presentation and other aspects of *in vivo* immunogenicity for live, attenuated vaccines such as BCG.

## MATERIALS AND METHODS

### Strains and media

For all experiments except Figures 5D-F, we used *M. tuberculosis* mc^2^6206 (H37Rv Δ*panCD* Δ*leuCD* (52)), in some cases transformed with pMV261-mEmerald for imaging (provided by Drs. Chris Sassetti and Christina Baer, University of Massachusetts Medical School). *M. tuberculosis* strains were cultured using Middlebrook 7H9 (liquid) or 7H10 (agar) supplemented with 10% Middlebrook Oleic Albumin Dextrose Catalase (OADC), 0.5% glycerol, 0.05% Tween 80, and kanamycin (20 μg/ml) or hygromycin (50 μg/ml) when necessary. Because *M. tuberculosis* mc^2^6206 is a double auxotroph, media were additionally supplemented with 50 μg/ml L-leucine (Sigma) and 24 µg/ml pantothenic acid (Sigma). We omitted these supplements for the experiment in Figures 5D-F, in which we used the wild-type parental H37Rv strain of *M.tuberculosis*.

Human monocyte/macrophage cell line THP-1 and immortalized bone marrow derived macrophages (iBMDM) from C57BL/6J mice were respectively gifts of Drs. Barbara Osbourne, University of Massachusetts Amherst, and Christopher Sassetti, University of Massachusetts Medical School (37). *VAMP2* knockout and wild-type THP-1 cells were purchased from Ubigene Biosciences, Guangzhou Science City, Guangdong, China (Cat # YKO-H1108). THP-1s were maintained in suspension in RPMI 1640 medium (Genesee Scientific) supplemented with 10% (v/v) heat-inactivated fetal bovine serum (FBS; Fisher Scientific), antibiotic-antimycotic (1% containing penicillin G, streptomycin, and amphotericin B), 1 mM sodium pyruvate, and 10 mM HEPES (all from Genesee Scientific) and maintained in a humidified incubator at 37 °C with 5% CO_2_ in air. iBMDMs were maintained in Dulbecco’s Modified Eagle Medium (DMEM; Genesee Scientific) supplemented with 10% FBS, 2 mM L-glutamine, and 10 mM HEPES and maintained in a humidified incubator at 37 °C with 5% CO_2_ in air. THP-1s were differentiated into macrophages by incubating cells with 200 nM, phorbol-12-myristate 13-acetate (PMA; Sigma) for 48 hr.

### Protein photocrosslinking and analysis

Stock solutions of synthetic O-x-AlkTDM, O-x-AlkTMM, and O-x-Alk-fatty acid (Figure 1C) were prepared in anhydrous DMSO at concentrations of 100 mM and stored at −20 °C. Other reagent stocks used in typical copper-catalyzed alkyne-azide cycloaddition (CuAAc) reactions included sodium ascorbate (60 mM in H_2_O, always freshly prepared); TBTA ligand (Click Chemistry Tools; 6.4 mM in DMSO; stored at –20 °C); CuSO_4_ (50 mM in H_2_O, stored at room temperature (RT)); TAMRA biotin azide (AzTB, Click Chemistry Tools, 10 mM in DMSO, stored at −20 °C).

Probes were diluted to 100 µM in DMSO or DMEM and added directly to the cells in solution or used to coat 100-mm cell culture dishes. Coated dishes were dried in a biosafety cabinet overnight. 5×10^6^ iBMDMs or differentiated THP-1s/plate were seeded onto the coated dishes and incubated for indicated time. Alternatively, macrophages were incubated +/- probe in solution for indicated period of time. Probe-treated and untreated control macrophages were then UV-irradiated (15-watt, 365 nm UV lamp) for 10 min on ice. Cells were washed with cold PBS and lysed with Pierce RIPA buffer (1 mM PMSF, 1x complete EDTA free protease inhibitor; Thermo Fisher). To reduce capture of endogenous biotinylated proteins, 500 µl of protein lysate was pre-cleaned using 50 µl of Pierce NeutrAvidin Agarose (Thermo Fisher) that was initially washed twice with phosphate-buffered saline (PBS) and twice with RIPA lysis buffer. Beads and lysates were incubated overnight with constant rotation at 4 °C. Beads were then centrifuged at 3000 x g for 3 min and the protein concentrations of the supernatants were quantitated using Pierce bicinchoninic acid (BCA) assay (Thermo Fisher). Prior to CuAAc, protein concentrations were normalized using RIPA buffer. Lysates were clicked with AzTB for 2 hr with constant shaking at 37 °C using the following typical CuAAC reaction on 187.5 µl of lysate with indicated final concentrations: 1 mM copper(II) sulfate (4 µl), 100 µM TBTA (3.12 µl), 1 mM sodium ascorbate (3.4 µl), and 100 µM AzTB (2 µl) to give a final volume of 200 µl. Proteins were precipitated using chloroform/methanol (1:1 v/v) at 4 °C to remove excess fluorophore and lipids. Precipitated protein pellets were resuspended in 200 µl of RIPA lysis buffer and incubated at 37 °C for 10 min for protein solubilization. 15 µl of each sample was saved as pre-enrichment click input. The remaining 185 µl of clicked lysate was incubated with 50 µl of Neutravidin beads that were pre-washed as described above. Beads were washed five times with RIPA buffer (200 µl) and five times with PBS (200 µl) via a series of centrifugation steps at 3000 x g for 3 min.

Biotinylated proteins were eluted by adding 30 µl of 4x Laemmli loading sample buffer (BioRad) and boiled at 95 °C for 15 min. ∼10-15 µg of input sample (click lysates) and eluted proteins were resolved by SDS-PAGE and analyzed by in-gel fluorescence. The rhodamine channel was used to detect TAMRA-labeled proteins and Coomassie staining was used to detect total protein (Cytiva; Amersham ImageQuant™ 800 Western blot imaging system).

For immunoblotting, 10 μg of the input controls and 15 μL of eluted proteins were resolved by 12% SDS-PAGE. After gel electrophoresis, the proteins were semi-dry transferred onto an Immun-Blot PVDF membrane (Thermo Fisher). PVDF membrane and filter paper were incubated in a transfer buffer (25 mM Tris, 193 mM glycine, and 20% methanol) and put into a transfer cassette for transfer using the TransBlot Turbo Transfer System. The time of protein transfer was based on protein size, ∼7-30 min, for smaller and larger proteins, respectively (25 V). After transfer, the membrane was blocked for 1 hr at room temperature in 5% dry nonfat milk in Tris-buffered saline containing Tween 20, pH 7.4 (50 mM Tris, 0.5 M NaCl, 0.01% Tween 20 (TBST)). Antibodies for immunoblotting were diluted 1:1000 in TBST containing 5% milk.

Primary antibodies included: rabbit polyclonal anti-Syntaxin 8 (1:1000), rabbit polyclonal antiserum for VTI1B (1:1000), and mouse monoclonal anti-Synaptobrevin2/VAMP2 (1:1000) from Synaptic Systems, Germany. Mouse monoclonal anti-Mincle (1:1000) from MBL International Corporation, rabbit polyclonal antiserum for Endobrevin/VAMP8 (1:1000 in 5% Bovine Serum Albumin (BSA) in TBST; Synaptic Systems), rabbit polyclonal anti-Syntaxin 5 (1:1000; Cell Signaling), rabbit monoclonal anti-Syntaxin 6 (1:1000; Cell Signaling), rabbit polyclonal anti-Syntaxin 18 (1:1000; Proteintech), rabbit monoclonal anti-GAPDH (1:1000) from Cell Signaling. Anti-rabbit IgG and anti-mouse IgG Horseradish peroxidase (HRP)-linked antibody (1:1000; Cell Signaling). The PVDF membrane was incubated overnight at 4 °C with constant shaking with primary antibodies. The membrane was then washed with 1x TBST thrice for 10 min. Secondary antibodies (anti-mouse IgG, and anti-rabbit IgG from Cell Signaling) were diluted 1:1000 in TBST containing 5% milk and the membrane was incubated for 1 hr at room temperature with constant shaking. PVDF membrane was then washed thrice with 1x TBST for 10 min and viewed with the Amersham ImageQuant™ 800.

### LC–MS/MS analysis

10 μL of samples were run into a BioRad Criterion TGX Precast Gel for ∼20 min at 50 V constant so that the samples just entered into the stacking gel and did not resolve. Electrophoresis was then stopped and the gel was stained using Coomassie Blue. The resulting protein bands were excised from the gel and subjected to in-gel tryptic digestion according to Trypsin gold manufacturing protocol (Promega) with modifications. Gel bands were destained with several incubations with 100 mM of ammonium bicarbonate (AB) and 1:1 (v/v) 50 mM ammonium bicarbonate/50% acetonitrile (ACN). Briefly, gel bands were dehydrated using 100% ACN and incubated with 10 mM dithiothreitol in 100 mM AB at 50 °C for 30 min, dehydrated again, and incubated in the dark with 55 mM iodoacetamide in 100 mM AB for 20 min. Gel bands were then washed with AB and dehydrated again. Trypsin gold was prepared to 20 μg/μL in 40 mM AB/10% ACN and added to each gel band so that the gel was completely submerged. Bands were then incubated at 37 °C overnight. Peptides were extracted from the gel by sequential 5 minute incubations with 1 % formic acid/2% ACN, 1:1 (v/v) ACN/water and 1% formic acid in ACN. Pooled supernatants were vacuum-dried and resuspended in 0.1% formic acid for ZipTip cleanup (C_18_ column; Sigma). Samples were then sent to the University of Massachusetts Amherst Mass Spectrometry Core where LC-MS analysis was performed using a Easy-nLC 1000 nanoLC chromatography system interfaced to an Orbitrap Fusion mass spectrometer (Thermo Scientific). Samples were pre-concentrated and desalted on a C18 trap column prior to separation over a 90-min gradient from 0% to 40% mobile phase B (A:0.1% formic acid in water, B:0.1% formic acid in ACN) at 300 nL/min flow rate with a 75 µm x 15 cm PepMap RLSC column (Thermo Scientific). Mass spectrometry parameters were as follows: ion spray voltage 2000V, survey scan MS1 120k resolution with a 2 s cycle time, interleaved with data-dependent ion trap MS/MS of highest intensity ions with HCD at 30% normalized collision energy. Raw files were analyzed in Proteome Discoverer 2.4 (Thermo Scientific) using the SEQUEST search algorithm using the *Mus musculus* Proteome database. The search parameters used were as follows: 10 ppm precursor ion tolerance and 0.4 Da fragment ion tolerance; up to two missed cleavages were allowed; dynamic modifications of methionine oxidation and N-terminal acetylation. Peptide matches were filtered to a protein false discovery rate of 5% using the Percolator algorithm. Resulting list of proteins were filtered for proteins that had at least 10% coverage and present in three out of the four biological replicates. Only proteins present in the O-x-AlkTDM-treated conditions and not in the controls were considered hits.

### Cytokine production

iBMDMs were incubated with indicated concentrations of O-x-AlkTDM or trehalose-6,6-dibehenate (TDB; InvivoGen) for indicated times. Supernatants were collected and TNF-alpha and IL-6 production were measured by enzyme-linked immunosorbent assay (R&D Systems, DuoSet ELISA).

### Coating green fluorescent latex beads with TDM

200 μl of non-functionalized green fluorescent latex beads (1 µm and 3 µm, Bangs Laboratories) were washed thrice with 1 ml of PBS. Beads were resuspended in 1 ml of 0.1 M borate buffer (Polysciences) and 100 µg Immunoglobulin G (IgG; Sigma), Bovine Serum Albumin (BSA; Fisher Scientific), O-x-AlkTDM, native TDM (Invivogen) and TDB (InvivoGen) were then added. The mixture was incubated overnight at RT with constant rotation. The beads were centrifuged for 10 min and resuspended in 1 ml of 10 mg/ml fat-free BSA (Sigma) in 0.1 M borate buffer, and incubated for 1 hr constant rotation. They were then centrifuged for 10 minutes, and the supernatant was discarded. The beads were washed thrice with PBS, then stored in 500 µl of PBS at 4°C in the dark until experimental use.

### Immunoprecipitations

#### Immunoprecipitation from M. tuberculosis-infected macrophages

iBMDMs were infected with *M. tuberculosis* mc^2^6206 for 4 hr at estimated multiplicities of infection (MOI; bacteria:macrophage) of ∼1 or ∼5. Because we have observed for *M. tuberculosis* that effective MOIs are 5-10% or less than intended MOI, we report estimated MOIs using the more conservative 10% conversion. After 4 hr, cells were washed thrice with L-leucine and pantothenate-supplemented DMEM and incubated an additional 6 hrs in the same medium. Macrophages were then lysed with RIPA buffer. Lysates were filtered twice through 0.2 μm filters (Corning Costar Spin-X; Sigma) and then pre-cleaned with protein G beads (Thermo Fisher) to remove non-specific binding. Protein concentration was quantified by BCA assay and 200 μg of protein was incubated with 2 μg/ml of primary antibody or with isotype control (rabbit IgG or mouse IgG isotype controls; Cell Signaling). Protein-antibody complexes were incubated overnight with constant rotation at 4 °C. Protein G beads (50 μl) were added to the protein-antibody complexes and incubated for 4 hr with constant rotation at 4 °C. Beads were washed four times with RIPA buffer and 30 μl of 4x Laemmli loading sample buffer was added to the beads. Beads were boiled for 15 minutes at 95 °C, and supernatants were collected for immunoblotting. Immunoblotting was performed as described above in “**Protein photocrosslinking and analysis**” using anti-SNARE antibodies and Tidyblot (1:1000; BioRad) in 5% nonfat milk in TBST. Tidyblot was used as a secondary antibody to prevent detection of denatured antibodies.

For Figures 5D-F, differentiated THP-1 cells were infected with wild-type *M. tuberculosis* H37Rv at estimated MOIs of ∼1 or ∼3 for 4 hrs. After 4 hrs cells were washed thrice with DMEM and incubated an additional 4 hrs in the same medium. Macrophages were lysed and immunoprecipitated as described above.

#### Immunoprecipitation from bead-incubated macrophages

iBMDMs were incubated with BSA, IgG-, O-x-AlkTDM-, TDB or TDM-coated beads (1 µm) at an effective multiplicity of infectIon (MOI) of ∼5 (macrophage:beads) for 1 hr. Cells were then washed thrice with cold PBS and lysed with RIPA buffer. Lysates were prepared and immunoprecipitations carried out as described in “*Immunoprecipitation from M. tuberculosis-infected macrophages”*.

#### Immunoprecipitation of TDM-SNAREs

As described in “**Protein photocrosslinking and analysis**”, iBMDMs were incubated with O-x-AlkTDM, UV-irradiated, lysed, and subjected to CuAAC with AzTB. Clicked lysates were then chloroform/methanol precipitated and resuspended pellets were immunoprecipitated with antibodies to VAMP2, VTI1B and STX8 at 2 μg/ml. To test for O-x-AlkTDM presence we used in-gel fluorescence (to detect TAMRA) and/or immunoblotted with anti-biotin antibodies (1:1000; 5% nonfat Milk in TBST; Abcam).

### eGFP-VAMP2 expression in iBMDMs

Lonza nucleofector with SG Cell Line 4D-Nucleofector™ X Kit S was used to generate eGFP-VAMP2-expressing macrophages. 2×10^5^ iBMDMs were washed and resuspended in the appropriate electroporation buffer. iBMDM cell suspensions were mixed with 2 µg of pEGFP VAMP2 (mouse; Addgene plasmid # 42308 (53). Cells were electroporated using the program FF-100 and immediately transferred to pre-warmed DMEM. Cells were plated in a 24-well plate for recovery, and after 48 hours, were sorted for eGFP-positive cells using BD FACSAria Fusion. Appropriate selection antibiotic was added after 24 hrs of sorting. After 6 days of antibiotic selection, cells were ready for experimental usage. Lysates were immunoblotted using anti-GFP (1:1000 in 5% non-fat milk in 1xTBST; Proteintech) and anti-VAMP2 to test for eGFP-VAMP2 expression (SI, Figure S7). We detected the fusion protein in iBMDMs that were successfully nucleofected with the plasmid and additionally observed that VAMP2 alone is more abundant in these cells.

### Macrophage infections

#### M. tuberculosis replication

1 ml stock of *M. tuberculosis* mc^2^6206 (previously frozen at OD ∼ 1.0) was thawed and diluted 1:10 in 7H9 medium and incubated at 37 °C with shaking at 150 rpm for 2-3 days, until the optical density at 600 nm (OD_600_) reached 0.3. One day prior to infection, iBMDMs were plated in a 96-well plate at a density of 2 × 10⁴ cells per well. THP-1 cells were plated at a density of 4 x 10^4^ cells and differentiated 48 hr before infection. On the day of infection, the bacterial culture was washed twice with PBS, and the OD_600_ was measured. The total *M. tuberculosis* required for the desired MOI was calculated based on this OD_600_ reading, and was transferred to DMEM or RPMI 1640 (depending on cell type) supplemented with 50 μg/ml L-leucine and 24 µg/ml pantothenic acid (without antibiotics). Medium was removed from the iBMDM or THP-1 plates, and fresh medium with resuspended *M. tuberculosis* was added at an estimated MOI of ∼1 (macrophages:bacteria), with 200 µl per well. Cells were incubated at 37 °C with 5% CO₂ for 4 hours. Afterward, the cells were washed thrice with medium, and fresh complete DMEM or RPMI 1640 was added to maintain continuous culture at 37 °C with 5% CO₂. For colony-forming unit (CFU) plating at 0, 24, 72 hr, and 120 hrs post-infection, macrophages were lysed with 0.1% Triton X-100 for 5 min, and the lysate was serially diluted using 0.05% Tween-80 in PBS. The diluted lysate was plated on 7H10 agar plates and incubated at 37 °C for 3-4 weeks. Colonies were counted, and CFUs per well were calculated.

#### Phagosome-lysosome fusion assay

Differentiated THP-1 cells were plated at 5×10⁴ cells per well in an 8-well Millicell EZ slide (Sigma) and then infected with mEmerald-expressing *M. tuberculosis* mc^2^6206 at an estimated MOI of ∼1 for 4 hrs. 100 nM Lysotracker (Thermo Fisher) was added for the last hour of infection and extracellular bacteria were removed by washing 3-4 times with cold PBS. For bead-containing phagosome-lysosome fusion assays, cells were incubated with 100 nM LysoTracker and 3 µm beads coated with BSA, TDB, O-x-alkTDM or TDM-coated beads at an estimated MOI of ∼1 for 1 hr. Non-internalized beads were then removed with 3-4 washes with cold PBS. Cells infected with *M. tuberculosis* or exposed to coated beads were fixed with 4% paraformaldehyde (Fisher Scientific) at RT for 10 minutes for bead-treated samples and 1 hour for *M. tuberculosis*-infected samples; washed thrice with PBS. 30 µl of VECTASHIELD® (Vector Laboratories) with DAPI was added to each well. The slide was mounted with a coverslip and sealed with nail polish. Fusion of bead or *M. tuberculosis*-containing phagosomes with lysosomes was quantified by blinded assessment of colocalization of mEmerald with Lysotracker using a Nikon TiE A1R-SIMe confocal microscope.

#### VAMP2 spatial distribution

iBMDMs expressing eGFP-VAMP2 were plated at 5×10⁴ cells per well in an 8-well Millicell EZ slide (Sigma) then infected with mCherry-expressing *M. tuberculosis* mc^2^6206 at an estimated MOI of ∼1 (macrophages:bacteria) for 4 hrs. As described in “*Phagosome-lysosome fusion assay*”, extracellular bacteria were washed and iBMDMs were fixed with 4% paraformaldehyde. Slides were mounted using VECTASHIELD® with DAPI. Images of infected and uninfected macrophages were acquired using a Nikon TiE A1R-SIMe confocal microscope.

#### Mycolic acid remodeling by O-TMM-C16

Stock solutions of synthetic O-TMM-C16 (80) were prepared in anhydrous DMSO at 25 mM and stored at −20°C. mEmerald-expressing *M. tuberculosis* mc^2^6206 was incubated with 1 mM O-TMM-C16 or 4 % final volume of neat DMSO for 5 days, with bacterial growth monitored daily via optical density (OD_600_) measurements. On day 5, differentiated THP-1 cells were infected with O-TMM-C16-labeled *M. tuberculosis* at an effective MOI of ∼1 for the indicated times.

Following infection, extracellular bacteria were removed as described in the “*Phagosome-lysosome fusion assay*,” and cells were fixed with 4% paraformaldehyde. Slides were mounted with VECTASHIELD® containing DAPI. Phagosome-lysosome fusion was quantified by blinded analysis of mEmerald colocalization with LysoTracker using a Nikon TiE A1R-SIMe confocal microscope.

#### Subcellular localization of O-x-AlkTDM

iBMDMs were seeded at 5 × 10⁴ cells per well in an 8-well Millicell EZ slide (Sigma) and treated with O-x-AlkTDM for the indicated durations. Cells were fixed with 4% paraformaldehyde for 10 minutes, followed by three washes with PBS. Permeabilization was carried out using 0.05% Triton-X (Sigma) for 15 minutes, and blocking was performed with 3% BSA in PBS (Fisher Scientific) for 30 minutes. CuAAC was conducted using a freshly prepared master mix containing 1 mM CuSO₄, 128 μM TBTA, 1.2 mM sodium ascorbate, and 20 μM Picolyl Alexa 488 azide-fluorophore (Click Chemistry Tools, 10 mM in DMSO, stored at −20 °C). Samples were incubated in the reaction mix at room temperature in the dark for 30 minutes with shaking, followed by three washes with 1% BSA in PBS. Finally, cells were washed with PBS, and slides were mounted as previously described.

## Acknowledgements

We thank Ms. Emily Lopes for help with blinding and Drs. Bill Wickner, Shumin Tan, Hesper Rego, and Dan Hebert for valuable discussions. We also thank the directors of the University of Massachusetts Amherst Mass Spectrometry, Light Microscopy, and Flow Cytometry facilities Drs. Stephen Eyles, James Chambers, and Amy Burnside for their help and advice. These studies were supported by the National Institutes of Health R21AI163949 (M.S.S. and B.M.S.), R15AI117670 (B.M.S.), R01AI177653 (A.C.R.), National Research Service Award T32 GM139789 (K.G.H.), and the National Research Service Award T32 GM135096 (P.N.L.), and the National Science Foundation 2320737 and 2117338 (B.M.S.). Figures created with BioRender.com.

## Supplementary Materials and Methods

### General experimental for synthesis

Reagents and solvents were procured from commercial sources without further purification unless otherwise noted. Anhydrous solvents were obtained either commercially or from an alumina column solvent purification system. All reactions were carried out in oven-dried glassware under inert gas unless otherwise noted. Analytical thin-layer chromatography (TLC) was performed on glass-backed silica gel 60 Å plates (thickness 250 μm) and detected by charring with 5% H_2_SO_4_ in EtOH. Column chromatography was performed using flash-grade silica gel 32–63 μm (230–400 mesh). NMR spectra were obtained using Bruker Avance 500 instrument. Coupling constants (*J*) are reported in hertz (Hz) with chemical shifts in ppm (δ) referenced to solvent peaks, with the following splitting abbreviations: s = singlet, d = doublet, dd = doublet of doublets, ddd = doublet of doublet of doublets, dt = doublet of triplets, t = triplet, td = triplet of doublets, m = multiplet. High-resolution electrospray ionization (HR ESI) mass spectra were obtained using an Agilent 6545XT LC-ESI-MS QTOF instrument.

### Synthesis of O-x-AlkTDM

Synthesis of O-x-AlkTDM was adapted from a previously reported procedure.^1^ An oven-dried round-bottom flask was charged with *N*,*N*’-dicyclohexylcarbodiimide (DCC) (0.207 g, 1.00 mmol) and *N*,*N*-dimethylamino-4-pyridine (DMAP) (0.063 g, 0.516 mmol). After drying the reagents under high vacuum and placing the flask under an argon atmosphere, anhydrous CH_2_Cl_2_ (5 mL) was added, and the mixture was cooled to 0 °C. To the stirring solution was added 9-(3-pent-4-ynyl-3-H-diazirin-3-yl-nonanoic acid^1^ (0.201 g, 0.760 mmol dissolved in 0.5 mL anhydrous CH_2_Cl_2_) and the reaction was stirred for 30 min. This was followed by slow, dropwise addition of freshly prepared solution of 2, 3, 4, 2’, 3’, 4’-hexakis-*O*-(trimethylsilyl)-α,α-trehalose^1^ (0.384 g, 0.495 mmol) in anhydrous CH_2_Cl_2_ (5 mL). The reaction mixture was stirred and gradually allowed to warm to room temperature. After TLC (hexanes/ethyl acetate 4:1) showed generation of mono- and diester products in approximately equal amounts (approximately 12 h), the reaction was quenched by addition of excess CH_3_OH and concentrated by rotary evaporation. The concentrated mixture was purified by silica gel chromatography (4:1 hexanes/ethyl acetate containing 1% Et_3_N) to give diester intermediate as an off-white solid. The intermediate was dissolved in CH_3_OH (5 mL) and Dowex 50WX8-400H^+^ ion-exchange resin was added and stirred for 30 min at room temperature, after which TLC (CH_2_Cl_2_/CH_3_OH 4:1) indicated the reaction was complete. The resin was filtered off and the filtrate was concentrated by rotary evaporation and purified by silica gel chromatography (CH_2_Cl_2_/CH_3_OH 9:1) to give O-x-AlkTDM (0.092 g, 29% over two steps) as a white solid. ^1^H NMR (500 MHz, CD_3_OD, referenced to methanol peak at 3.34 ppm): δ 5.07 (d, *J* = 3.5 Hz, 2 H), 4.39 (dd, *J* = 1.6, 11.9 Hz, 2 H), 4.23 (dd *J* = 5.3, 11.9 Hz, 2 H), 4.04 (ddd, *J* = 1.5, 5.1, 10.0 Hz, 2 H), 3.81 (t, *J* = 9.3 Hz, 2 H), 3.50 (dd, *J* = 3.6, 9.7 Hz, 2 H), 3.36 (t, *J* = 9.4 Hz, 2 H), 2.37 (t, *J* = 7.2 Hz, 4 H), 2.24 (t, *J* = 2.7 Hz, 2 H), 2.18 (dt, *J* = 2.7, 6.9 Hz, 4 H), 1.65 (pent, *J* = 7.0 Hz, 4 H), 1.51 (t, *J* = 5.3 Hz, 2 H), 1.42-1.26 (m, 26 H) 1.16-1.08 (m, 4 H). ^13^C NMR (125 MHz, CD_3_OD, referenced to methanol peak at 49.86 ppm): δ 176.3, 96.0, 85.0, 75.4, 74.0, 72.8, 72.4, 71.0, 65.3, 35.9, 34.7, 33.6, 31.2, 31.11, 31.09, 30.97, 30.1, 26.9, 25.7, 24.9, 19.4. HRMS (ESI-TOF) m/z [M+Na]^+^ Calcd for C_42_H_66_N_4_O_13_Na 857.4520; found 857.4524. Note that the monoester product generated in the esterification step was also obtained and desilylated according to the reported procedure to give O-x-AlkTMM (41% over two steps), which matched literature-reported NMR data (34).

### Synthesis of O-x-AlkTMM and x-AlkFA

Synthesized according to literature-reported procedures (34).

### BSA photocrosslinking

The labeling procedure was performed on defatted, reduced, and alkylated bovine serum albumin (BSA) prepared as in previously reported procedures.^1^ Briefly, BSA was incubated with excess CHCl_3_ overnight with stirring. After 12 h, BSA was collected by filtration and dried under air. A 10 mg/mL stock solution in water was prepared and diluted to 3.8 mg/ml. Dithiothreitol (DTT) (100 mM stock in 50 mM ammonium bicarbonate solution) was added to a final concentration of 5 mM and the protein was incubated at 56 °C for 20 min in a thermomixer. To reduced BSA, iodoacetamide was added to a final concentration of 16.5 mM (from 550 mM iodoacetamide stock in 50 mM ammonium bicarbonate solution) and incubated at room temperature in the dark for 20 min. To 90 μL aliquots of reduced and alkylated BSA was added 10 μL of previously reported N-x-AlkTMM-C15^1^ as positive control or O-x-AlkTDM synthesized here to a final probe concentration of 100 μM, BSA concentration of 2.7 mg/mL, DMSO concentration of 10%. The mixture was incubated at room temperature with mixing for 1 h. The BSA-probe mixtures were split into two equal volumes in 1.5 mL centrifuge tubes. The samples were either exposed to UV irradiation for 15 min or not using a 15-watt 365 nm UV bench lamp (UVP) at a distance of ∼2 cm. All six samples were subjected to Cu-catalyze azide-alkyne cycloaddition (CuAAC) using carboxyrhodamine 110 azide (Click Chemistry Tools) (30 μM) in the presence of sodium ascorbate (1.2 mM), copper(II) sulfate (1 mM) and tris(benzyltriazolylmethyl)amine (TBTA) ligand (128 μM), giving a final BSA concentration of 1.95 mg/ml. The reactions were incubated for 1 h at room temperature. Excess CuAAC reagents were removed by sequential addition of 4 volumes of methanol, 1 volume of chloroform, and 3 volumes of water. Samples were briefly vortexed and centrifuged at 18,000 x g for 5 min. The top methanol layer was carefully aspirated to not to disturb protein wafer. 3 volumes of methanol were added, followed by brief vortex and centrifugation at 18,000 x g for 5 min followed by careful aspiration. The resulting protein pellets were dried for 10 min and resuspended in 100 μL water. 20 μg of each sample was analyzed by 4-20% SDS-polyacrylamide gel electrophoresis (BioRad) in a Tris-glycine-SDS running buffer, followed by fluorescence scanning using a Typhoon FLA 7000 (GE Healthcare Life Science) with a 485 excitation and 520 emission filter. The gel was fixed for 15 min (40% ethanol, 10% acetic acid in milliQ water), rinsed once with milliQ water and stained over night with gentle agitation in QC Colloidal Coomassie stain (Bio-Rad). The gel was rinsed with milliQ water until the background was clear and imaged using a ChemiDoc Touch Imaging System (Bio-Rad) and processed by Image Lab software (Bio-Rad).

## Supplementary Figures

**Figure S1.**
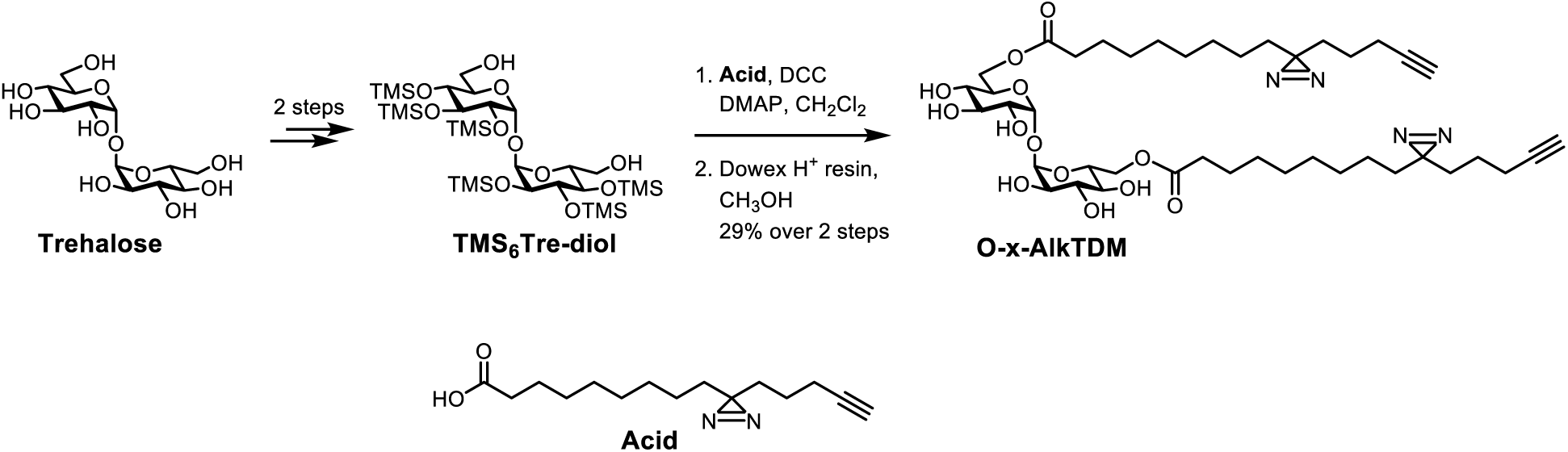
Synthesis of O-x-AlkTDM.

**Figure S2.**
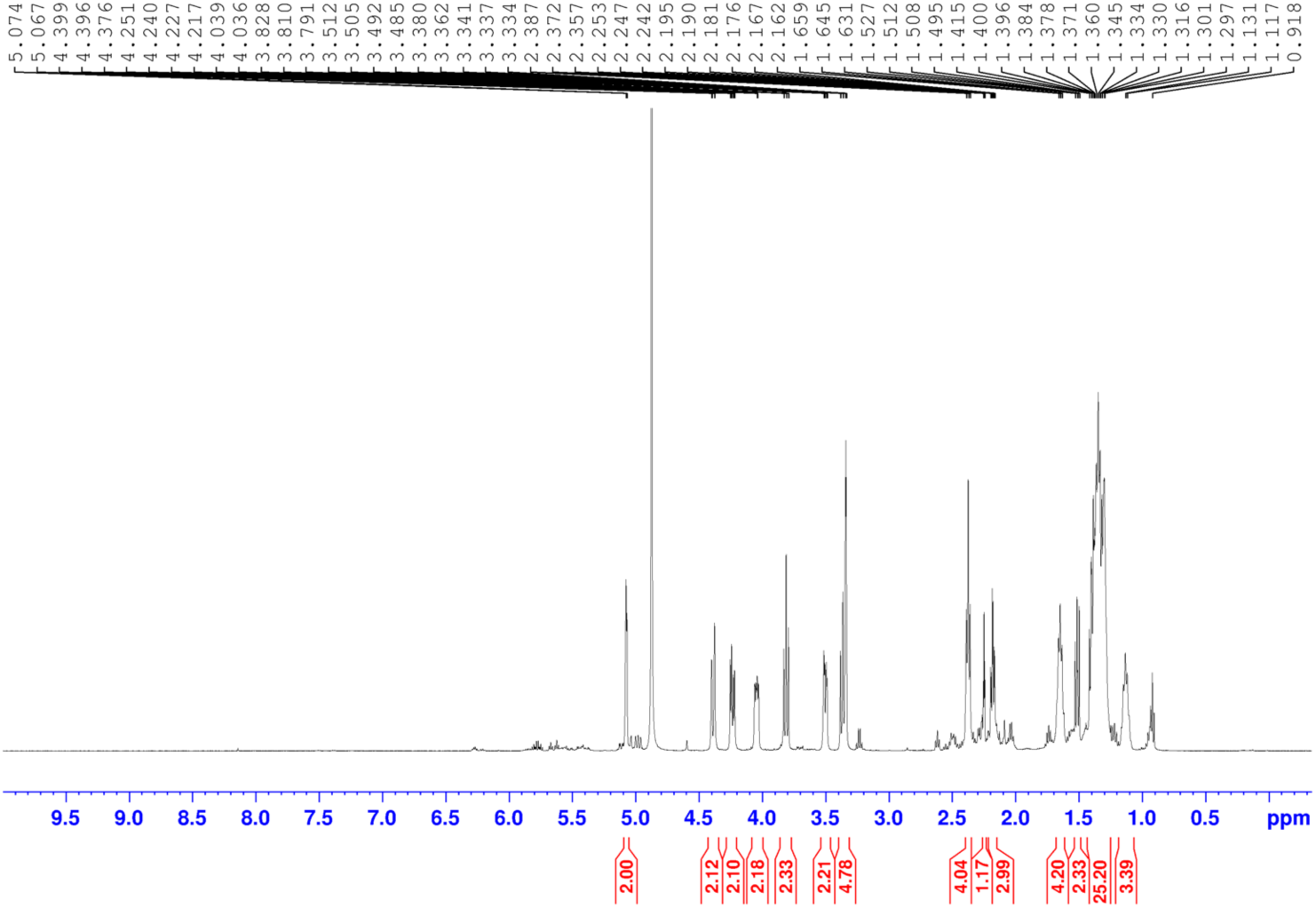
^1^H NMR spectrum of O-x-AlkTDM.

**Figure S3.**
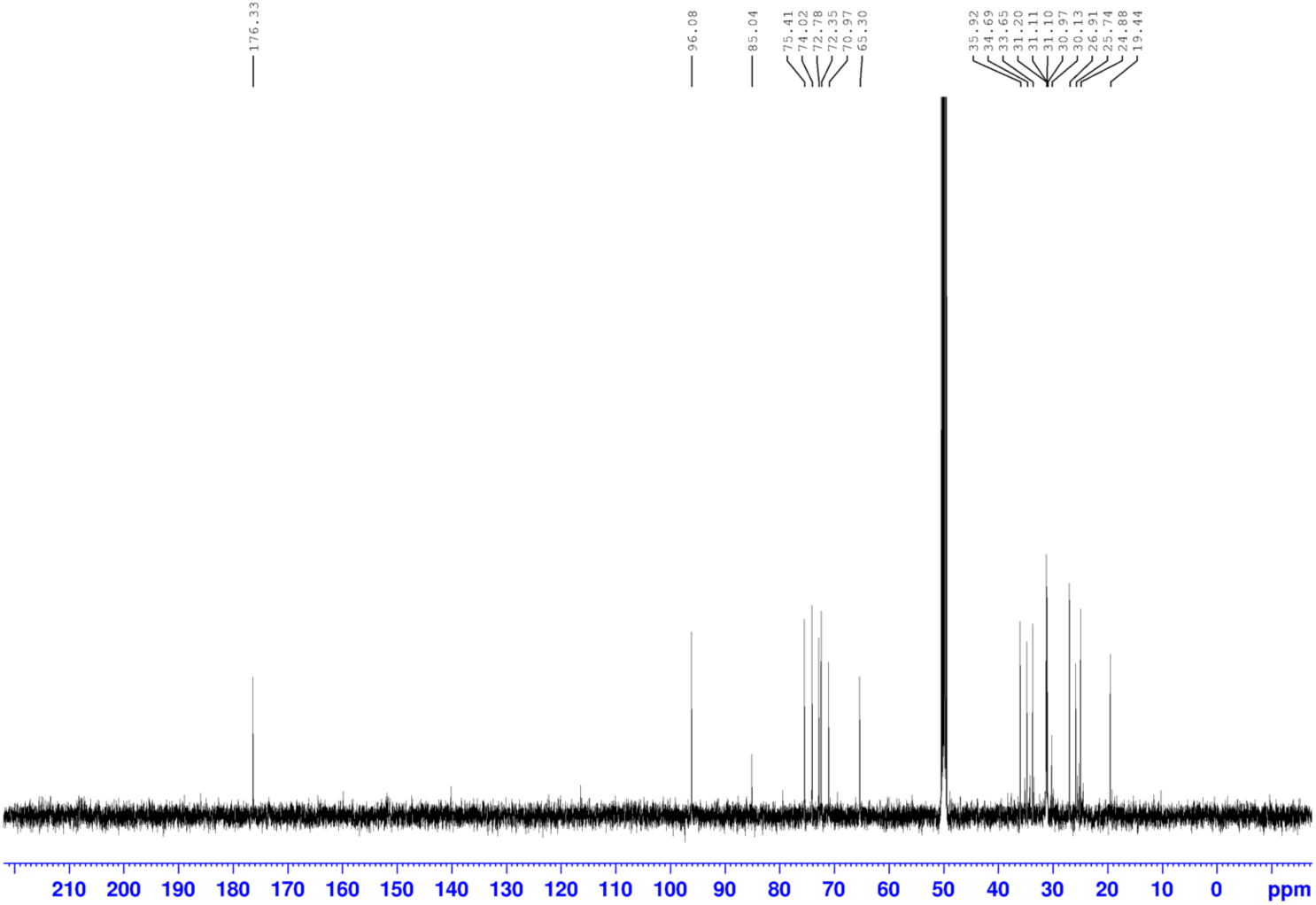
^13^C NMR spectrum of O-x-AlkTDM.

**Figure S4.**
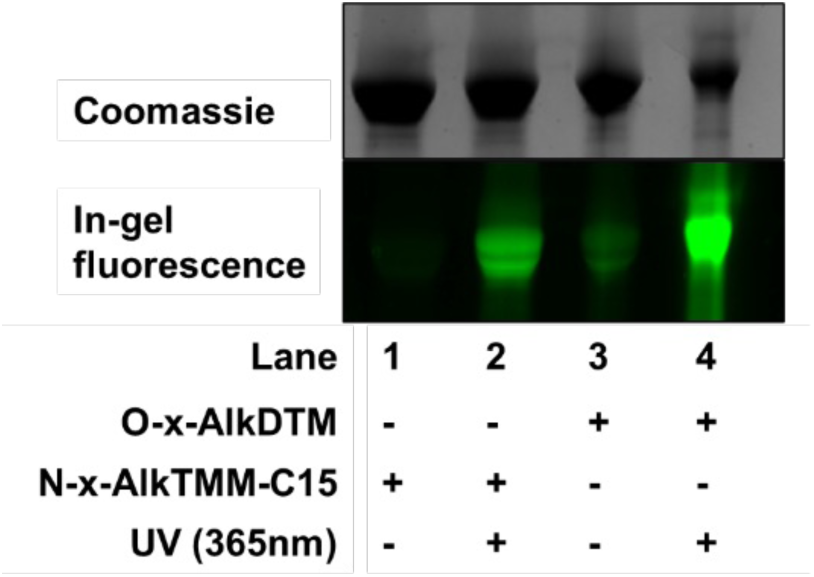
UV-dependent photo-crosslinking of bovine serum albumin (BSA) with O-x-AlkTDM (or positive control N-x-AlkTMM-C15) (34), followed by CuAAC-mediated fluorescence labeling and SDS-PAGE analysis with fluorescence scanning.

**Figure S5.**
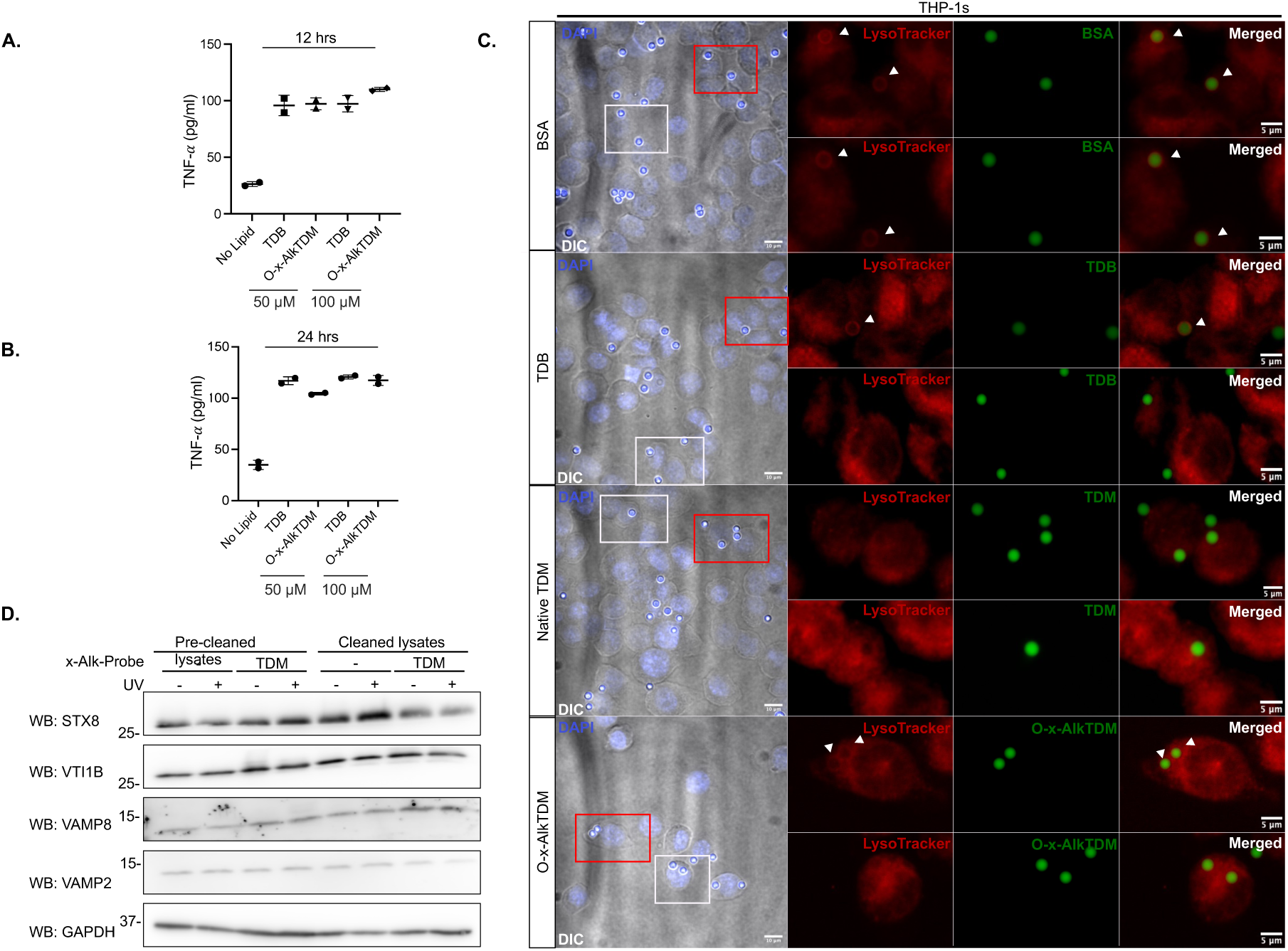
Supplement to. **Figure 2**. ELISA detection of pro-inflammatory cytokine TNF-alpha after 12 (A) or 24 (B) hour incubation of iBMDMs with O-x-AlkTDM or TDB. “No Lipid” represents unstimulated controls. Data are mean ± SD from two independent experiments. C) Additional images for experiments in Figure 2C. Lysosome fusion of bead-containing phagosomes. THP-1 cells were incubated with BSA-, TDB-, TDM-, or O-x-alkTDM-coated fluorescent beads (green), followed by LysoTracker staining to assess acidification. Quantification of spatial coincidence between beads and LysoTracker found in Figure 2D. White arrows indicate LysoTracker-localized beads. The red box highlights the top magnified image, while the white box corresponds to the bottom image for each condition. (D) Example of protein expression before and after clean up with NeutrAvidin agarose beads in iBMDMs treated with -/+ UV and O-x-Alk-TDM. Whole cell lysates were collected and analyzed via SDS-PAGE, followed by immunoblotting with indicated anti-SNAREs antibodies. There is no apparent difference between pre-cleaned and cleaned lysates prior to CuAAC with TAMRA Biotin Azide (AzTB).

**Figure S6.**
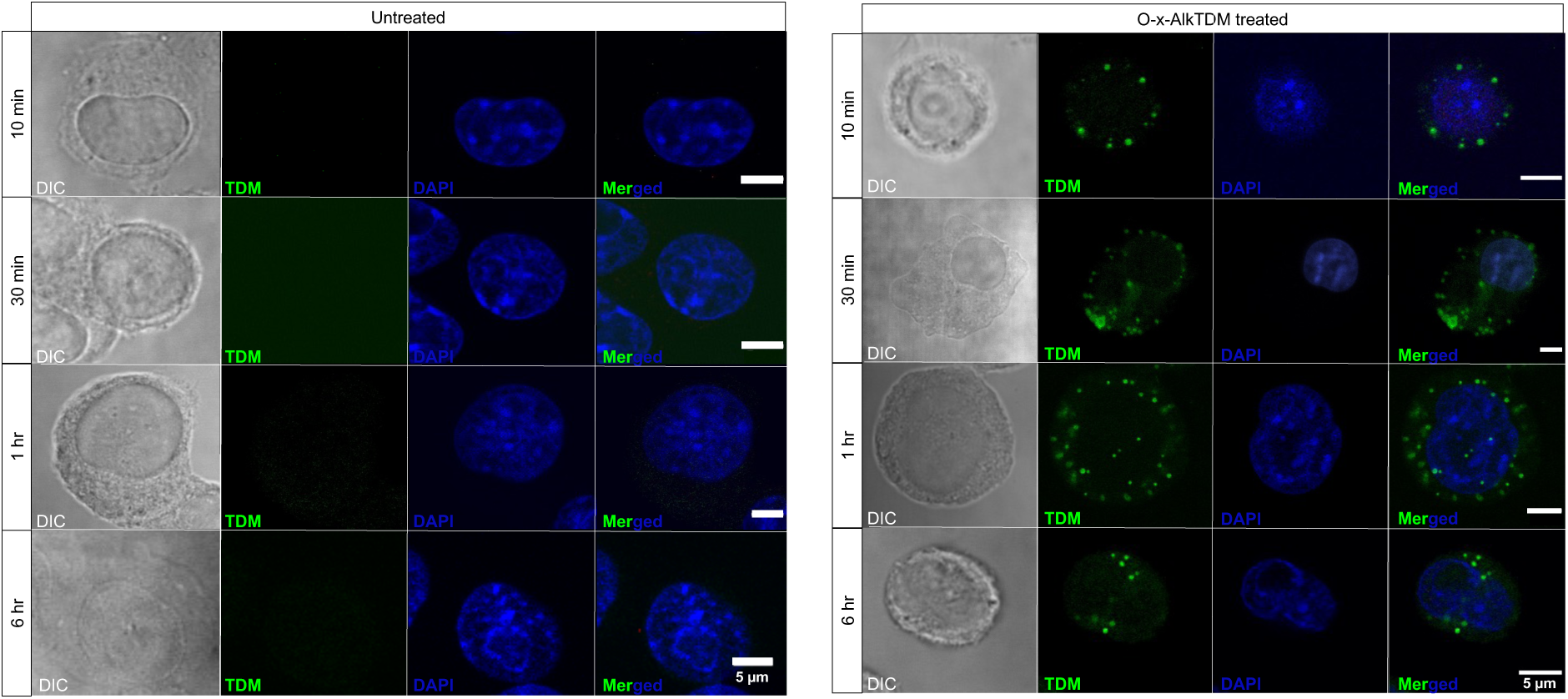
**Subcellular distribution of O-x-AlkTDM in iBMDMs**. Representative confocal microscopy images of iBMDMs in the presence of the absence (left) or presence (right) of the O-x-AlkTDM probe (green). iBMDMs were incubated +/- O-x-AlkTDM for the indicated time points, fixed, and subjected to CuAAC with Picolyl Alexa 488 Azide. Green fluorescent puncta appear to be in the macrophage cytoplasm. Differential interference contrast (DIC) images depict the overall cellular morphology and DAPI stains the nuclei.

**Figure S7.**
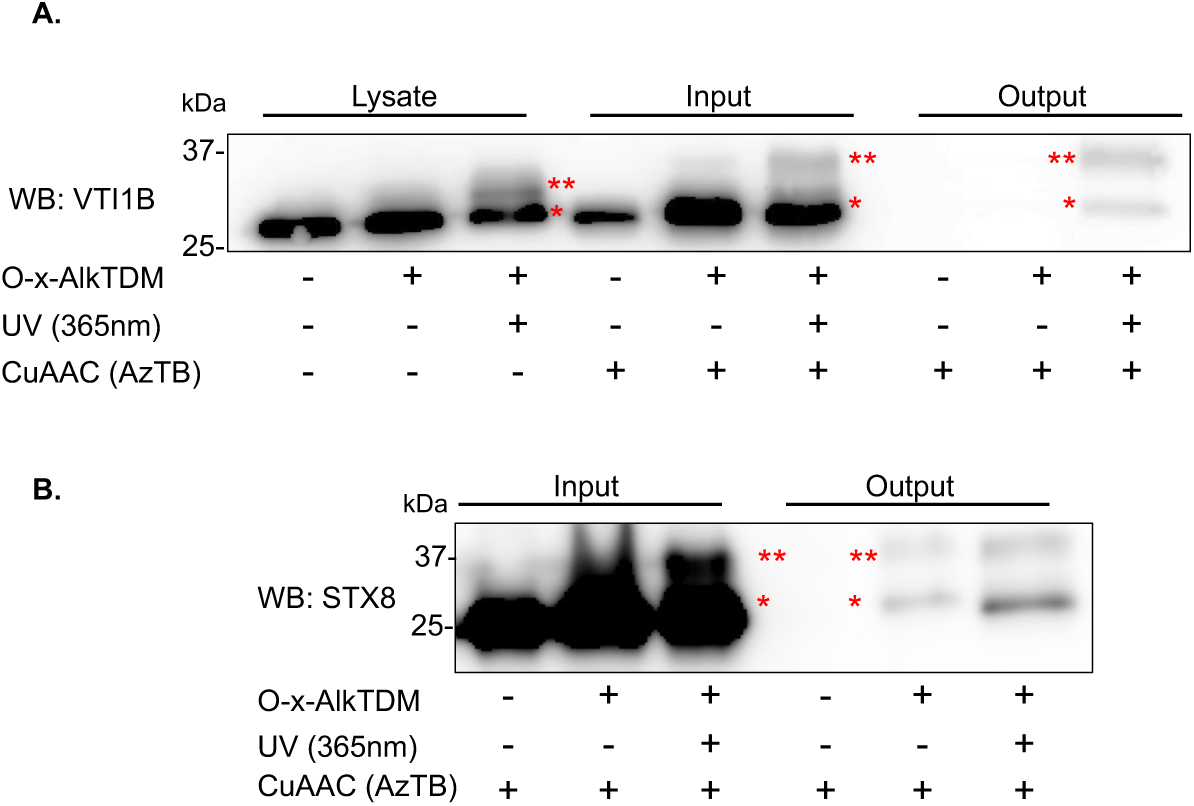
**VTI1B and STX8 may interact with more than one molecule of O-x-AlkTDM**. iBMDMs were treated 6 hrs +/- O-x-AlkTDM and processed as in Figure 2E-F. Samples were immunoblotted with the indicated antibodies prior to CuAAC (lysates), after CuAAC but prior to affinity purification (input), or after both manipulations (output). The presence of multiple bands (denoted with one or two asterisks), and their apparent increase in size after CuAAC (A), suggest the possibility of one or two O-x-AlkTDM molecules binding per SNARE.

**Figure S8.**
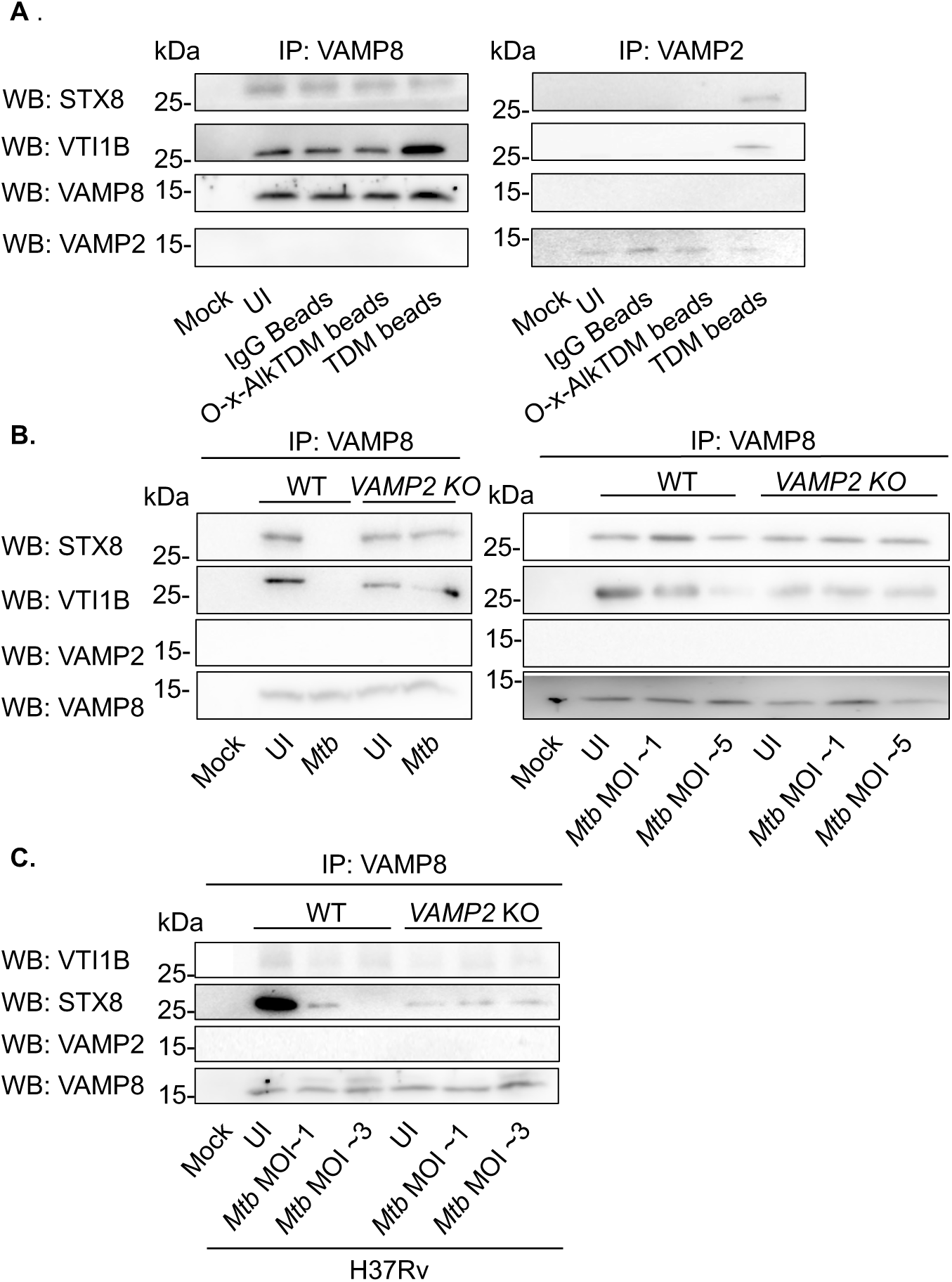
Independent biological replicates of experiments in. **Figures 4C, 4F and 5A-B.** (A) iBMDMs were incubated +/- latex beads coated with IgG, O-x-AlkTDM, or native TDM then processed as in Figures 4C and **4F**. Figures 4D and **4G** show quantitation of these experiments plus additional biological replicates from Figures 4C and **4F**. Wild-type (WT) and *VAMP2* KO THP-1s infected +/- *M. tuberculosis* mc^2^6206 (*Mtb*; estimated MOI ∼5, left, or estimated MOI ∼1 and ∼5, right) in (B) or +/- fully-virulent *M. tuberculosis H37Rv* (estimated MOI ∼1 or ∼3) in (C) and immunoblotted with anti-SNAREs after immunoprecipitation. In both strains, VAMP8 complexation with VTI1B and STX8 with high MOI *M. tuberculosis* is rescued in the absence of VAMP2. Mock, isotype control for each primary antibody. Figures 5B and **5E** show quantification of these experiments plus additional biological replicate from Figures 5A and **5D**, respectively. UI, untreated (A) or uninfected (B-C).

**Figure S9.**
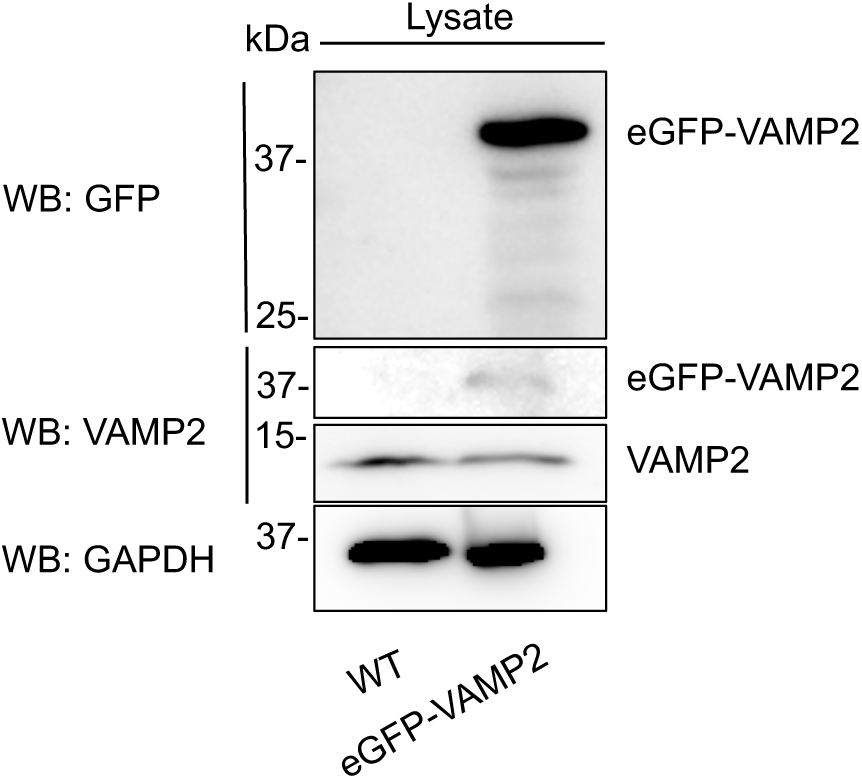
eGFP-VAMP2 expression in iBMDMs. Lysates (20 µg/lane) from wild-type (WT) and eGFP-VAMP2-nucleofected iBMDMs were immunoblotted for GFP, VAMP2, and housekeeping protein GAPDH. A band at ∼40 kDa (eGFP is 27 kDA and VAMP2 is 13 kDa) was detected in nucleofected iBMDMs by both antibodies.

**Figure S10.**
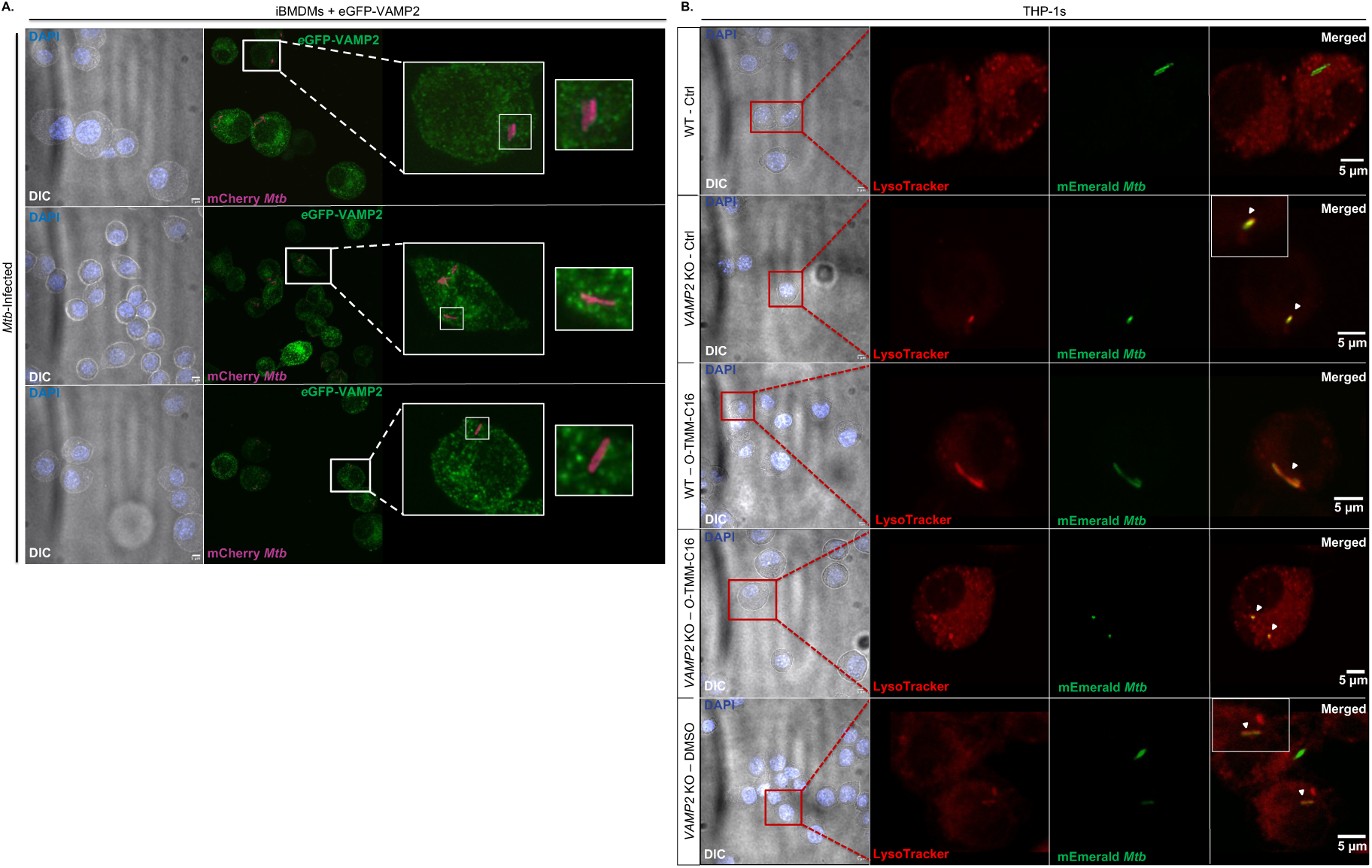
Additional images for experiments in. Figures 5G-H. (A) iBMDMs expressing eGFP-VAMP2 were infected with or without mCherry-expressing *M. tuberculosis* (estimated MOI ∼1) as in Figure 5G. (B) THP-1 cells infected with mEmerald-expressing *M. tuberculosis* (estimated MOI ∼1) were analyzed for spatial coincidence with LysoTracker as in Figure 5H.

**Figure S11.**
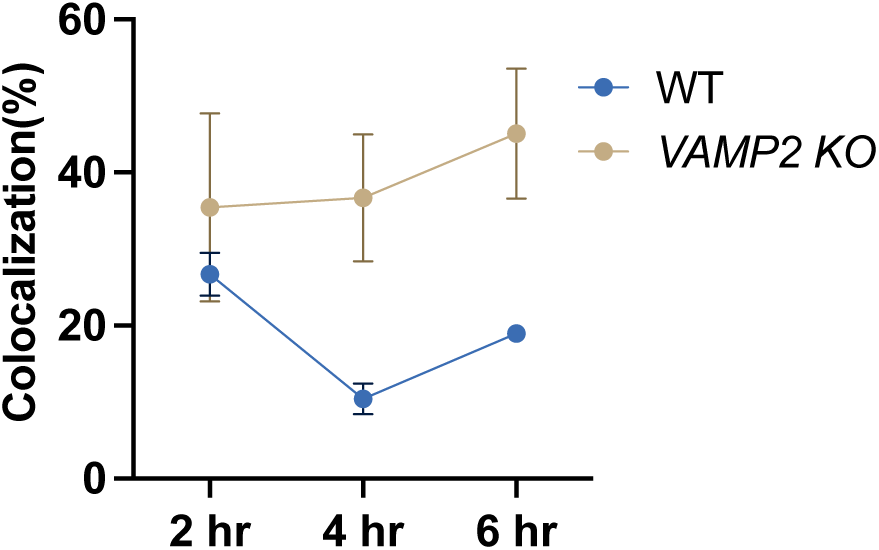
Phagosome acidification over time. Wild-type (WT) and *VAMP2* knockout (KO) THP-1 cells were infected with mEmerald*-*expressing *M. tuberculosis* and spatial coincidence with LysoTracker was assessed at the indicated time points. Data are mean ± SD from 3-6 independent experiments. The 4 hr time point data are the same as in Figure 5H (Ctrl).

**Figure S12.**
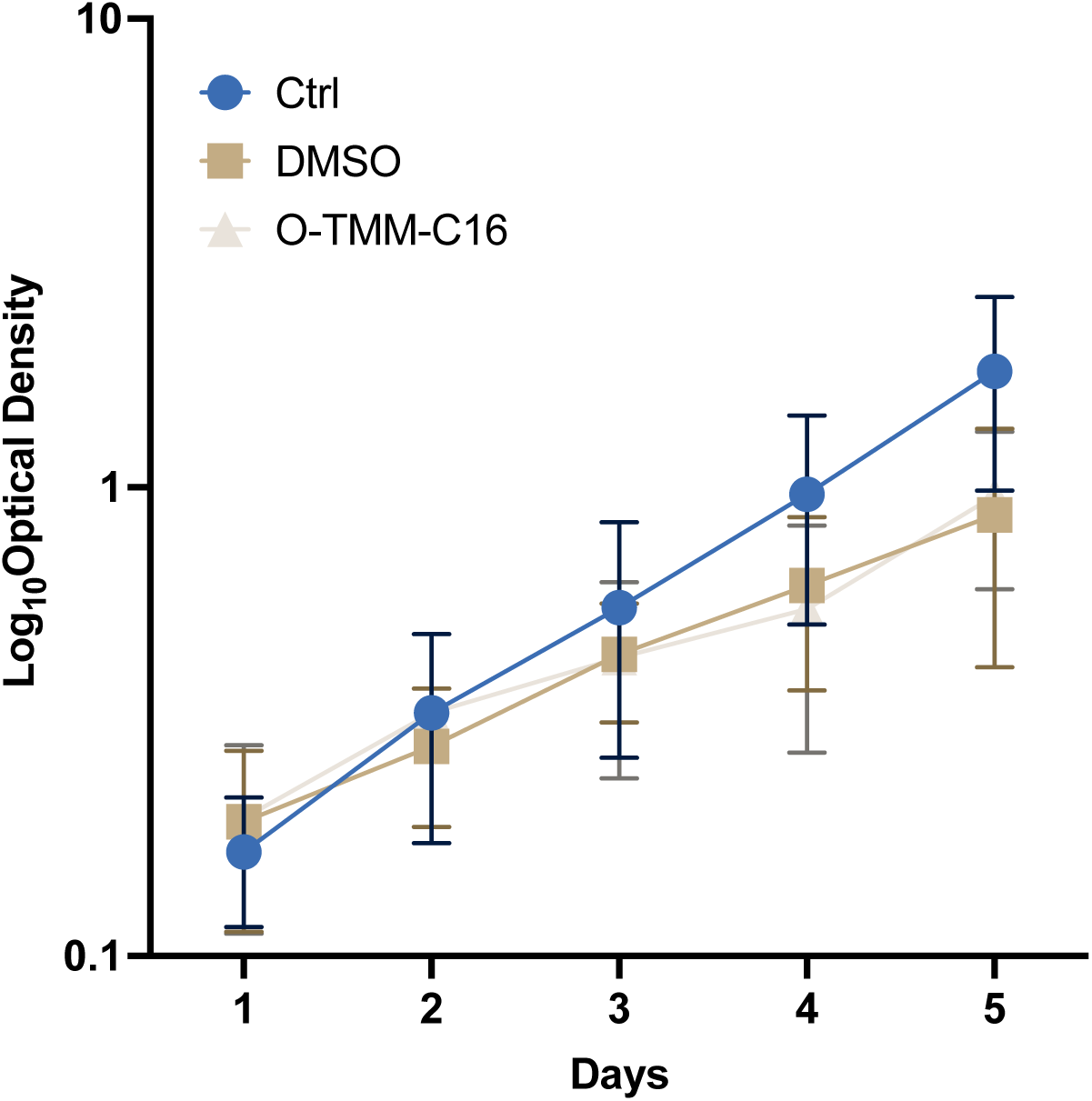
Growth of *M. tuberculosis* metabolically labeled or not with O-TMM-C16 (prior to macrophage infection in. Figure 5H). OD_600_ was monitored over 5 days for mEmerald-expressing M. tuberculosis incubated in 7H9 media alone (Ctrl); DMSO (DMSO); equivalent amount of DMSO and O-TMM-C16. Data are mean ± SD from three independent experiments

## Author Contributions

C.S., B.M.S., and M.S.S. designed research; C.S., K.J.B., P.N.L., J.C., C.Y.K., J.R.L., I.V.G., C.P., and K.G-H. performed research; C.S., B.M.S., and M.S.S. analyzed data; and C.S., B.M.S., and M.S.S. wrote the paper.

